# Hippocampal reactivation of aversive experience enables safety learning and slow-breathing state for recovery from stress

**DOI:** 10.1101/2025.06.03.657500

**Authors:** Baptiste Mahéo, Sophie Bagur, Dmitri Bryzgalov, Chloé Hayhurst, Mathilde Chouvaeff, Ella Callas, Carole Schmidt, Thierry Gallopin, Karim Benchenane

## Abstract

Adaptive threat responses require both defensive behaviours to minimize danger and recovering from the induced physiological stress. However, the behavioural and neural basis of these recuperative strategies are still elusive. Using a novel two-location fear conditioning paradigm in mice, we have identified a slow-breathing immobility state of recovery that emerges when animals identify safe environments after threat avoidance. This immobile state was characterized by a 2-4 Hz breathing profile and replay of the aversive experience in the hippocampus. Suppressing hippocampal sharp-wave ripples (SWRs) inhibited the emergence of this recovery state, suggesting their role in learning safe locations. Anxiolysis with diazepam directly promoted the recovery state while suppressing SWRs, showing this treatment to be a double-edged sword that facilitates immediate relief but impairs long-term safety learning. These results demonstrate the importance of hippocampal replay for emotional resilience through its role in recovery.

## Introduction

Animals face a dual challenge when confronted with threats such as predation or conspecific aggression. First, they deploy a cascade of defensive behaviours to avoid capture, such as freezing, avoidance, escape, or attack^1,2^. These strategies are essential to avoid physical harm but induce physiological and psychological stress that disrupts normal behaviours, including feeding, resting, and mating^3,4^. This leads to the second challenge: recovering from the stress induced by these events to restore homeostasis^5^. The ability to recover varies widely among individuals, and deficiencies can lead to long-term pathological consequences^6^. However, recuperation is currently ill-defined and therefore understudied. Relief from stress promotes rewarding neural processes that predict long-term resilience^7^ but have not been associated with a specific behaviour. Sleep has been investigated as a recovery-promoting condition^8^ and displacement behaviours such as self-grooming have been suggested as a potential coping strategy, yet their direct roles remain untested^9,10^. Therefore, whether animals engage in active processes for stress recovery remains unknown.

Focusing solely on behavioural markers may overlook critical aspects of the aspects of the recovery process, such as autonomic states, which are increasingly recognized as active participants in coping with threat. For instance, 4 Hz breathing actively contributes to the regulation of freezing behaviour^11^; while slow breathing can be artificially induced in mice to reduce stress^12,13^ and is widely used as a relaxation technique in humans^14^. Given this active role, breathing emerges as a potentially powerful physiological marker for distinguishing between defensive and recuperative states.

The hippocampus has been proposed as a key structure in stress and anxiety regulation^15,16^, a function eclipsed by its role in spatial learning and memory^17^. However, a key challenge in aversive learning is extracting and integrating threat-related information while minimizing direct exposure to danger. The hippocampus, via awake replay of past or possible events during sharp-wave ripples (SWRs), enables vicarious exploration and planning^18^, making it a strong candidate for this process. Supporting this idea, hippocampal awake replay of aversive events has been correlated with threat avoidance^16,17^. This could allow animals to steer clear of danger without needing to encounter it directly, yet its causal role remains untested. Beyond avoiding threat, effective stress recovery also requires accurately determining when and where it is safe to engage in recuperative behaviours. This requires distal representations of threat that could be facilitated by replay. This perspective suggests that the hippocampus not only helps avoid threats but also identifies safe zones for recovery. Here we used a spatial avoidance task to expose animals to threat while providing a safe zone to assess the emergence of recovery behaviours that could rely on hippocampus replay given the spatial nature of the task. With this approach, our results identify a slow-breathing recovery state that actively reduces stress levels and relies on hippocampal replay-driven identification of the safe zone.

### Identification of two different immobility states associated with distinct breathing signatures

In the U-Maze spatial avoidance task, mice were conditioned to avoid an eyelid shock when entering a randomly assigned “shock arm” while the other “safe arm” was never associated with shock (Fig. 1a). To minimize anxiety related to novelty, mice were habituated to the U-Maze one day prior to conditioning. On the conditioning day, we first recorded baseline behaviour and sleep patterns in the home cage (Sleep Pre) (Fig. 1a). We then assessed arm preference across four pre-conditioning sessions, confirming equal initial exploration of both arms (Fig. 1b). Mice then underwent 9–12 conditioning sessions, receiving a shock 13 ± 1 s after entering the shock arm. As expected, mice quickly learnt to avoid the shock zone (Fig. 1c). Because mice rapidly escaped upon stimulation, in a subset of sessions, we temporarily blocked them in either the shock or safe arm to allow more detailed behavioural sampling in these zones. Following conditioning, a second home cage sleep session assessed task-induced changes (Sleep Post). Finally, in a shock-free test phase in the U-Maze, we evaluated post-conditioning behaviour and observed strong avoidance of the shock arm (Fig. 1b).

**Figure 1.**
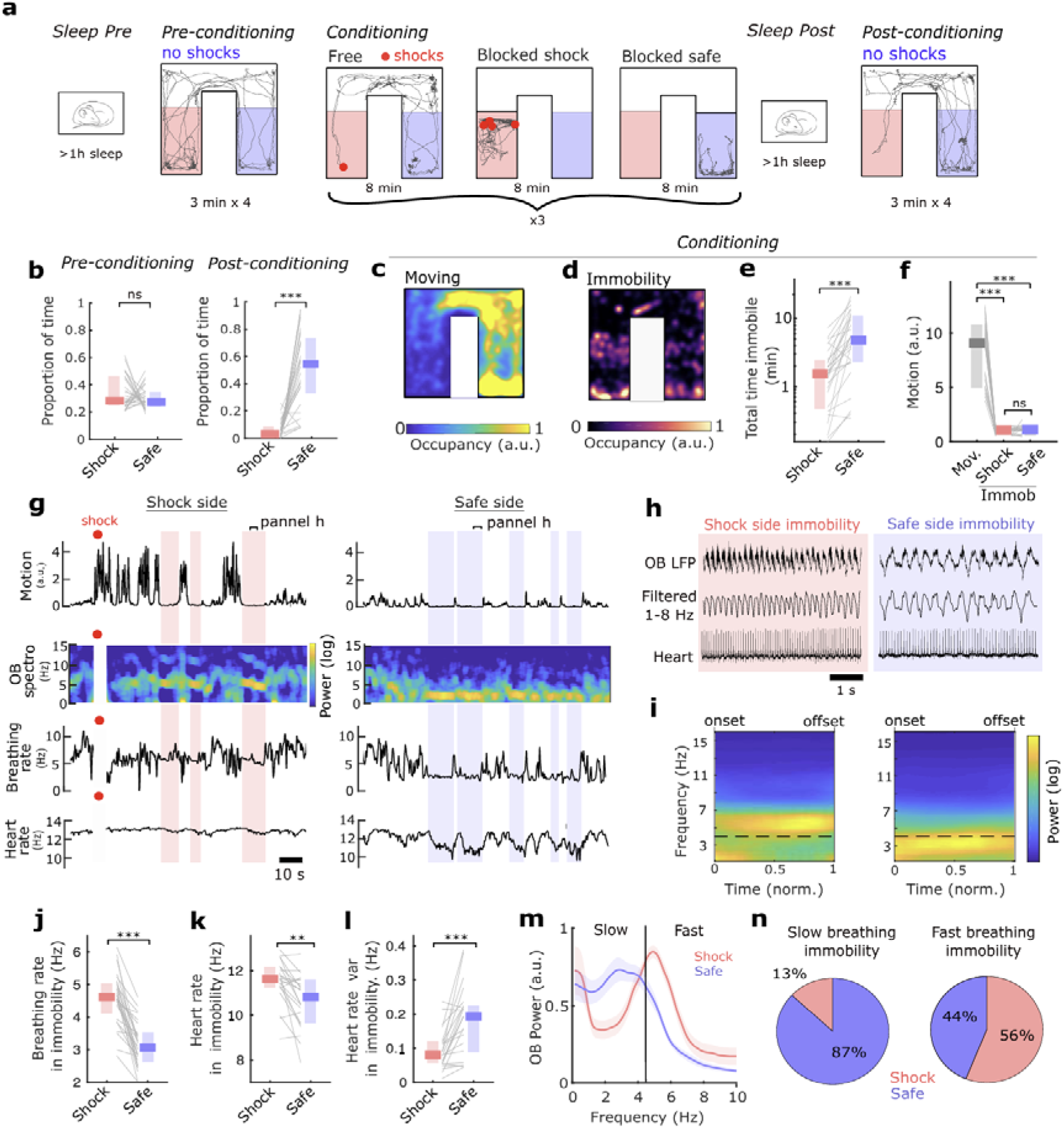
Identification of slow and fast breathing immobility in shock and safe arms. a. Schematic of U-Maze protocol with example trajectories and shock locations for one mouse. b. Proportion of time spent in shock and safe arm during the pre- and post-conditioning sessions (n=29, (NS)P=0.3695, ***P=5.32 **×** 10^−6^, Wilcoxon signed rank test). c. Average occupancy map during unblocked conditioning sessions when the mouse is actively exploring (n=29). d. Average occupancy map during conditioning sessions, including blocked and unblocked, when the mouse is immobile (n=29). e. Total time spent immobile in shock or safe arm during conditioning (n=28, ***P=9.78 **×** 10^−6^, Wilcoxon signed rank). f. Mean head acceleration (motion) during locomotion and immobility in shock or safe arm. (n=28, ***P=2.70 **×** 10^−5^ and 1.82 **×** 10^−5^, (NS)P=0.93, Wilcoxon signed rank test). g. Head acceleration (motion), OB spectrogram, respiratory rate and heart rate during time periods displaying both exploration and immobility in both arms for an example mouse. Note the clear difference in all physiological parameters during immobility in each arm. h. Raw data from shock and safe side immobility from bracketed time points in g. i. Mean OB spectrogram from all immobility periods in shock and safe arm, after normalizing duration between 0 and 1 (n=28) The yellow band corresponds to the breathing oscillation that shows the clear shift in frequency from 4-6 to 2-4 Hz between arms. j-l. Mean breathing rate (j), heart rate (k), and heart rate variability (l) during the final 10% of immobility in shock and safe arm epochs (***P=4.72 **×** 10^−6^, 2.01 **×** 10^−4^ and 3.34 **×** 10^−4^, Wilcoxon signed rank test). m. Mean OB spectrum during shock and safe arm immobility. The black line shows the breathing rate limit (4.5 Hz) used to distinguish slow and fast breathing immobility (n=28, error bars are s.e.m.). n. Proportion of slow (left) and fast (right) breathing immobility periods observed in the shock or safe arms during conditioning (n=28) All box plots show median and quartiles. Detailed statistics are provided in supplementary table 1.

During conditioning sessions, in addition to the avoidance behaviour, mice exhibited prolonged periods of complete immobility throughout the U-Maze, remaining immobile for an average of 1.6 minutes in the shock arm versus 5.2 minutes in the safe arm (Fig. 1d,e), corresponding to approximately 10% of the time spent in each arm. The quantity of head motion during immobility was indistinguishable between the two arms (Fig. 1f), and based on all standard metrics used to assess fear responses, these episodes would typically be classified as ‘freezing’ (Fig. S1a).

We compared physiology during immobility in both arms by recording local field potentials (LFPs) in the olfactory bulb (OB) to extract the respiratory rhythm (n=29) and electrocardiogram-derived heart activity (n=22). As is visible in the raw physiological traces (Fig. 1g,h, Supp. Video), shock arm immobility was characterized by 4-6 Hz breathing and safe arm immobility by 2-4 Hz breathing. Moreover, heart rate was higher and heart rate variability lower in the shock arm compared to the safe arm, indicating increased parasympathetic activation in the safe arm^21^. Additionally, tail temperature was lower during shock immobility, consistent with blood flow redistribution toward the limbs in preparation for flight^22^ (Fig. Sup1b) and gamma oscillations in the OB—markers of vigilance state^23^—were of higher amplitude and frequency during shock immobility (Fig. S1c-e). We trained a linear classifier to use these physiological variables to discriminate between immobility in the two arms (Fig. S1j). Cross-validating across individuals, it achieved an accuracy of 80%, and the classifier score, a continuous index reflecting the similarity of a given immobility period to shock- or safe-arm physiological profiles, demonstrated a clear distinction between conditions (Fig. S1k). This score will be used to quantify the global physiological state throughout the remainder of the manuscript.

These differences were robust across animals, especially late in conditioning, after mice had learned the task (Fig. 1i-l) and in all session types including blocked and unblocked, as well as later recall sessions without shock and in a separate group of mice using dorsal periaqueductal gray as a negative reinforcer (Fig. S2a-d). Moreover, a similar pattern emerged in a social defeat stress paradigm. When exposed to the larger, aggressive male, mice exhibited 4-6 Hz immobility in the aggressor’s cage whereas in their home cage, immobility was characterized by a lower 2–4 Hz frequency (Fig. S2e,f).

Constructing a linear model to predict breathing rate during immobility showed that the difference between shock and safe immobility types was largely explained by maze position and not by other factors (Fig. S3a-c). The distinction between the two breathing modes is not simply due to closeness to the painful stimulation since the breathing rate difference is observed at all post-shock latencies (Fig. S3a,b) and the intensity of aversive stimulation had no impact on breathing rate (Fig. S3h). Moreover, neither movement prior to immobility, time spent immobile nor overall time within the experimental protocol accounted for this difference (Fig. S3a-g). Thus, to assess the systematic relationship between physiological type and location in the maze, we categorized all immobility periods into slow (<4.5 Hz) and fast (>4.5 Hz) breathing modes (Fig. 1m). Fast breathing immobility was observed, as expected in the shock arm, but also in the safe arm in the early stage of the conditioning experiment. This pattern reflects learning, as discussed in detail below (Fig. 5). In contrast, slow breathing immobility was overwhelmingly expressed only in the safe zone (Fig. 1n), indicating that the absence of threat may be crucial for its emergence.

Together, these observations identify two distinct types of immobility with different physiological profiles occurring close and distant from threat respectively, raising their question of how they each contribute to dealing with threat.

### Slow breathing immobility is a recovery state distinct from freezing

We compared these two immobility profiles to the most widely studied defensive behaviour: freezing, evoked by context or sound after classical conditioning. Freezing, as previously reported^11,24^, was characterized by breathing above 4 Hz and its overall physiology resembled fast breathing immobility found in the shock arm (Fig. S4). We conclude that fast breathing immobility corresponds to freezing, a defensive strategy deployed by the animal in the presence of threat to avoid detection and prepare for flight^1,25^ (Fig. 2a).

**Figure 2.**
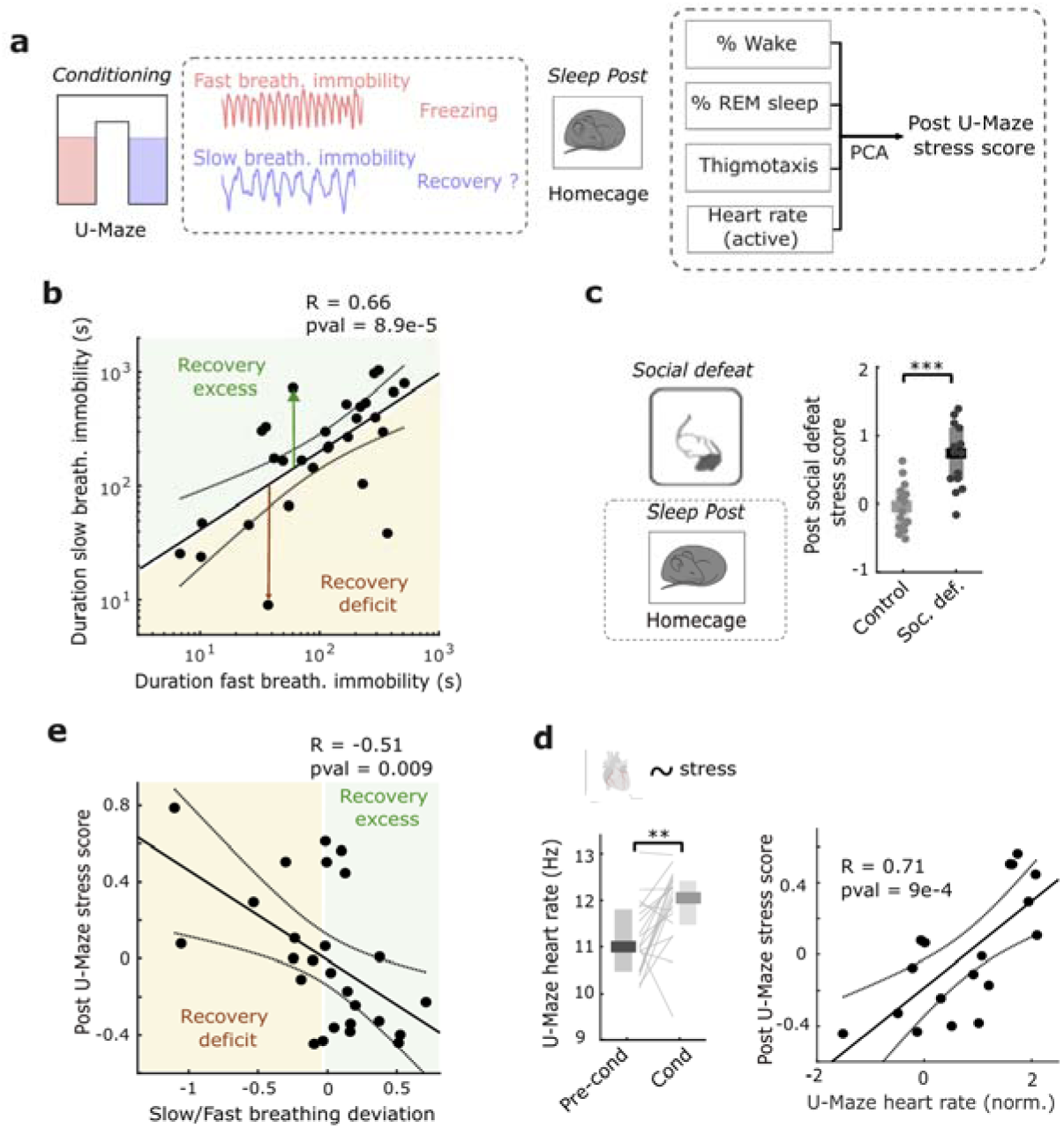
Slow breathing immobility predicts post U-Maze recovery from stress. a. Schematic of proposed interpretation of fast and slow breathing immobility. Fast breathing immobility corresponds to previously identified defensive freezing and slow breathing immobility is postulated to correspond to a recovery state. This is tested by relating it to a post U-Maze stress score that combines four classical stress measures in the homecage: insomnia, REM sleep loss, thigmotaxis and heart rate. b. Duration of fast breathing immobility state is strongly correlated with duration of slow breathing immobility state. This allows to define the slow/fast breathing deviation for each mouse (arrows) as the excess or deficit of slow breathing immobility duration relative to that expected based on the regression over all mice (n=28, R=0.67, ***P=8.90 × 10-5, Pearson correlation). c. After a social defeat stress, the stress score is increased compared to non-stressed controls (n=14 sessions, ***P=4 × 10-4, Wilcoxon rank-sum test). d. Heart rate during locomotion in the U-Maze task shows a significant increase during the conditioning phase (n=20, ***P=2.5631 × 10DD, Wilcoxon signed rank test), and is subsequently correlated with the post U-Maze stress score (n=20, R=0.71, ***P=9.27 × 10DD, Pearson correlation). Heart rate is normalized for each mouse to a baseline period in the homecage to reduce inter-individual variability e. The slow breathing deviation defined in d is correlated with the post U-Maze stress score (n=26, R=-0.50632, **P=0.00901, Pearson correlation). All box plots show median and quartiles. Detailed statistics are provided in supplementary table 1.

Slow breathing immobility on the other hand is distinct from freezing (Fig. S4b-d). Characterized by slow breathing and parasympathetic cardiac signatures (Fig. 1j-l), it suggests a calm state that may facilitate stress recovery. This is supported by a striking correlation between the duration of fast- and slow-breathing immobility (Fig. 2b). If fast breathing immobility, corresponding to freezing, is indeed a stressful defensive state and slow breathing immobility a putative recovery state, this relationship suggests that the greater the stress, the longer the compensatory recovery period.

In this case, deviation from this balance should predict individual animals’ resistance to the stress induced by the task. We therefore measured stress after conditioning with four well-established metrics: increased thigmotaxis and heart rate during wakefulness, global insomnia and reduced REM sleep proportion^26–28^. These features were all correlated so we applied PCA to derive a global stress score with almost identical contributions of each parameter (Fig. 2a, Fig. S6a). This score, and the individual stress markers, were validated by their increase in mice exposed to social defeat stress (Fig. 2c, Fig. S6b). Moreover, heart rate during locomotion is increased by conditioning in the U-Maze (Fig. 2d) even when corrected from locomotor activity (Fig. S6e). This increased heart rate during the task correlated positively with the post-task stress score, thus relating stress during and after the U-Maze task (Fig. 2d).

We quantified for each mouse how much slow-breathing immobility duration deviated from the expected duration based on the linear relation with the time the animals spent in fast-breathing immobility (“slow/fast breathing deviation”) and related it to the post-task stress score (Fig. 2e). This shows that mice with an excess of slow-breathing immobility during the task tended to have lower post-task stress scores, whereas those with a deficit exhibited elevated stress scores. These results hold also for individual stress parameters (Fig. S6c). The stress score was not correlated with overall time immobile (Fig. 6d). Moreover, breathing frequency during safe immobility episodes was positively correlated with the stress score (Fig. S7a) and all individual stress metrics (Fig. S7b). Thus, the slower the breathing, the lower the mouse’s post-task stress levels. In contrast, shock immobility breathing rate showed no correlation (Fig. S7b). Importantly, slow breathing immobility is physiologically distinct from quiet wake and sleep (Fig. S5) and does not occur prior to aversive stimulation (Fig. S3d), indicating that it is an actively regulated response to threatening situations.

These findings support slow breathing immobility as a mechanism for stress recovery. Moreover, they reveal a dynamic balance between stress accumulation—reflected in the fast breathing freezing state—and its compensatory reduction through the slow breathing putative recovery state.

### Hippocampal Reactivation of Aversive Experiences During Recovery

Slow breathing immobility is specific to the safe zone of the U-Maze (Fig. 1n) and therefore its expression likely depends on spatially identifying the shock-free zone in the environment. We therefore investigated hippocampal activity occurring during this state. We recorded LFP activity from the hippocampus (n=29) and simultaneously monitored single-unit activity in a subset of animals (n=9 mice, 484 single units) (Fig. 3a). This revealed a strong shift in hippocampal activity from low-frequency theta oscillations during fast breathing freezing to irregular activity with a significantly higher incidence of SWRs during slow breathing immobility (Fig. 3b, Fig. S1h,i). This transition is classically associated with a shift from externally driven sensory processing to internally generated representations^29^. In addition, SWR occurrence correlated with slower breathing frequency (Fig. S8a).

**Figure 3.**
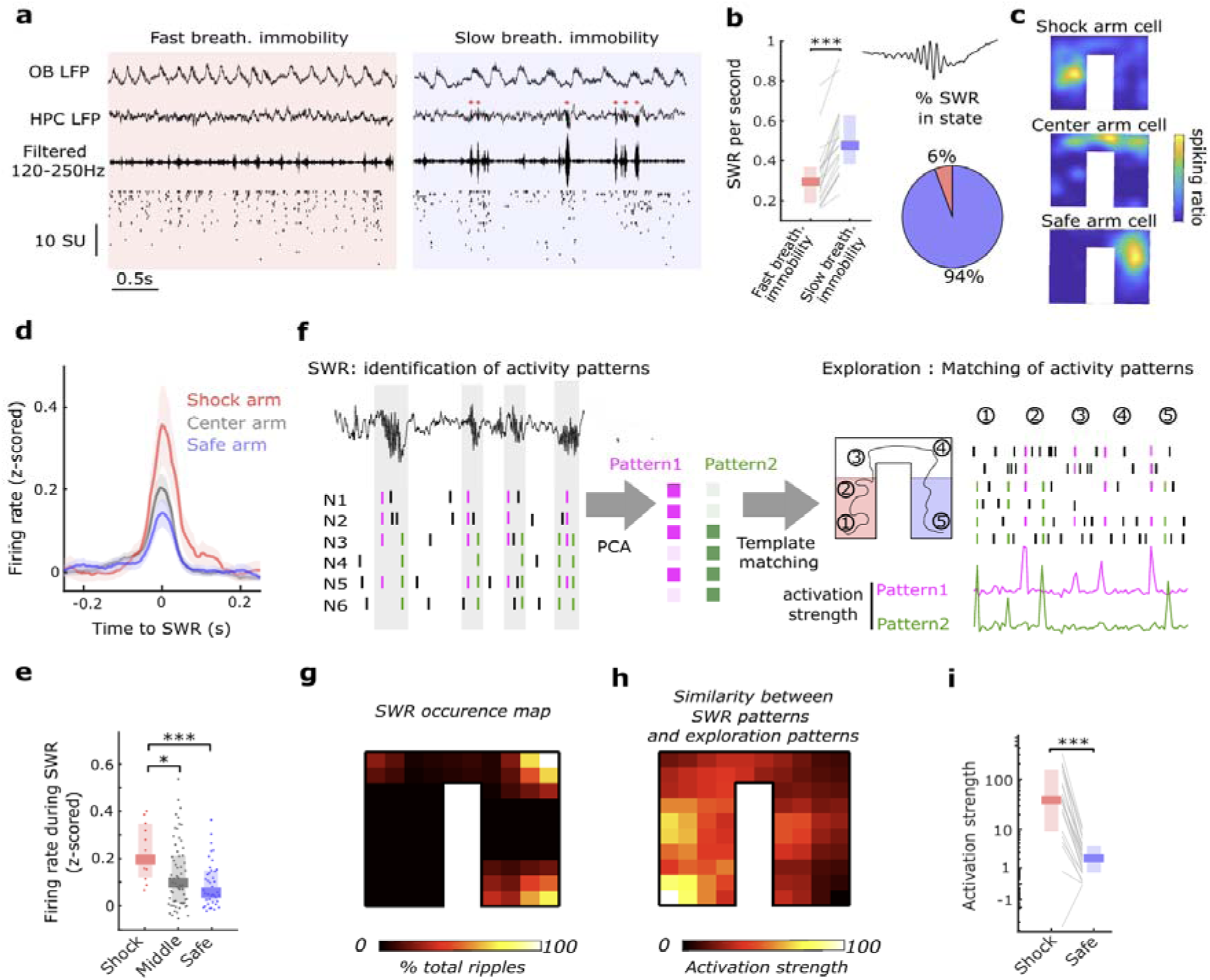
Slow breathing immobility is characterized by numerous HPC SWRs and reactivation of shock arm activity. a. Raw data showing OB and HPC LFP, HPC LFP filtered in the ripples frequency band and single unit activity from HPC single units during fast and slow breathing immobility. b. Ripples per second is increased in the slow breathing immobility state relative to the fast breathing immobility (n=19, ***P=1.32 × 10-4, Wilcoxon signed rank test). c. Firing maps of three example place cells during conditioning in the U-Maze. d. Ripple-triggered firing of place cells grouped according to the location of their place field : in the shock arm, in the center arm and in the safe arm. (n=159 place cells, error bars are s.e.m). e. Mean firing during the 50 ms around ripple peak for place cells grouped according to the location of the place field showing that shock arm place cells are more strongly activated (n=21, *P=0.012, ***P=5.34 × 10-4, Wilcoxon rank-sum test). f. Illustration of the method for identifying population patterns of activity during ripples using PCA. These patterns are then matched to data during exploration to measure their instantaneous activation strength. This allows calculation of where these patterns tend to occur in the U-Maze. g. Spatial map of ripple occurrence (n=9). h. Spatially averaged activation strength for patterns identified during ripples (n=22 patterns) i. Mean activation strength of all ripple-identified patterns during shock or safe side exploration showing that these patterns more strongly correspond to activity in the shock arm (n=22 patterns, ***P=1.32 × 10-4, Wilcoxon signed rank test) All box plots show median and quartiles. Detailed statistics are provided in supplementary table 1

Given the well-established role of SWRs in the reactivation of prior experiences, we investigated the information contained in neural activity during SWRs. During the conditioning phase, we recorded n=159 place cells with place fields distributed throughout the maze (Fig. 3c). All cells exhibited increased firing during SWRs. Those cells with place field locations in the shock zone were most strongly activated during SWRs (Fig. 3d,e), suggesting that mice were preferentially reactivating the aversive experience during these events.

To verify this at the population level, we identified recurring patterns of coordinated neuronal activity—proxies for cell assemblies—activated during SWRs using PCA. We then compared the similarity of these activity patterns to those observed during active exploration, computing a reactivation strength index (Fig. 3f). Reactivation strength was highest for locations in the shock zone, despite SWRs overwhelmingly occurring in the safe-zone (Fig. 3g-i). This effect was specific to SWRs, as constructing reactivation templates using non-ripple periods of safe immobility did not yield the same link with the shock arm (Fig. S8b).

These analyses show that during slow breathing immobility, an elevated occurrence of SWRs supports the replay of aversive experiences.

### Hippocampal Sharp-Wave Ripple Disruption Prevents the Emergence of the Recovery State

We next asked whether SWRs played a role in promoting slow breathing immobility, the putative recovery state. To test this hypothesis, we inhibited hippocampal activity during SWRs using a closed-loop stimulation protocol targeting the ventral hippocampal commissure, previously used to demonstrate a causal role of SWRs in memory^30^. In the “ripple inhibition” group, stimulation was triggered upon SWR detection during immobility throughout the conditioning protocol (Fig. 4a), with an average delay of 12 ms, 75% specificity, and 78% selectivity (Fig. S8c,d)^31,32^. To control for nonspecific effects of stimulation, a sham group received the same electrical stimulations following a delay (200±50 ms) after SWRs detection (Fig. 4a). Both groups received identical numbers of stimulations (Fig. S8e,f).

**Figure 4.**
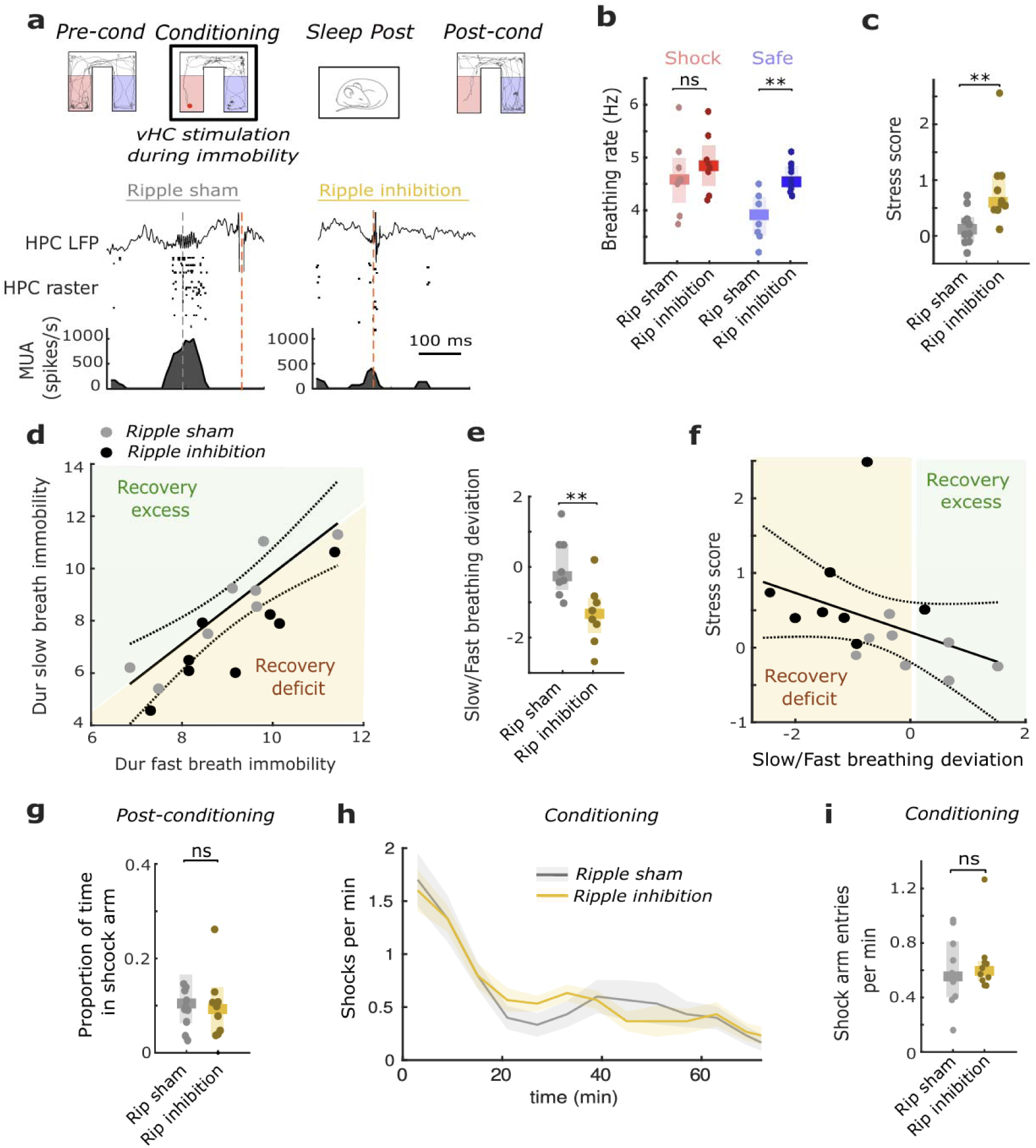
SWR disruption reduces slow breathing immobility and impairs stress recovery. a. Scheme of protocol showing vHC stimulation during conditioning sessions. Example data shows an online detected ripple that triggers a vHC stimulation that suppresses ongoing firing and an online detected ripple that received a sham, delayed stimulation b. Mean breathing rate in shock and safe arm for SWR disrupted and sham groups showing a specific increase in safe arm immobility breathing frequency (n=8/8, **P=0.0023, (NS)P=0.40, Wilcoxon signed rank test). c. Post U-Maze stress score for SWR disrupted and sham groups showing an increase in post-task stress after SWR disruption (n=10/10, **P=0.0046, Wilcoxon signed rank test) d. Time in the slow breathing immobility state is strongly correlated with time in the slow breathing immobility state for SWR disrupted and sham groups. SWR disrupted animals show a systematic shift towards lower amounts of slow breathing immobility, which we quantify with the slow/fast breathing deviation relative to the relationship defined using sham mice. Note that for this analysis, we excluded mice for which the instantaneous breathing rate could not be correctly estimated due to the stimulation artefact (sham, n=8, R=0.68, *P=0.049, inhib, n=8, R=0.72, *P=0.024, Spearman correlation). e. Slow breathing deviation is more strongly negative for SWR disrupted animals relative to sham controls (n=8/8, **P=0.0093, Wilcoxon rank-sum test). f. The slow breathing deviation is correlated with the post-task stress score across the two groups (n=16, R=-0.52, *P=0.037, Spearman correlation). g. SWR disruption does not affect time spent in the shock arm during post-conditioning test sessions relative to sham controls (n=10/10, (NS)P=0.73, Wilcoxon rank-sum test). h. SWR disruption does not affect the temporal evolution of shocks received per minute throughout the conditioning sessions relative to sham controls, showing no difference in learning dynamics (error bars are s.e.m.). i. SWR disruption does not affect the total number of shocks during the conditioning sessions relative to sham controls (n=10/10, (NS)P=0.73, Wilcoxon rank-sum test). All box plots show median and quartiles. Detailed statistics are provided in supplementary table 1

SWR disruption specifically modified slow-breathing immobility in the safe arm: breathing rate increased in the safe arm of the maze and the decoder classified safe side immobility as shock-like based on all physiological variables (Fig. 4b, Fig. S8g). However, SWR disruption had no effect on immobility in the shock zone.

Remarkably, SWR-disruption during the U-Maze conditioning led to an increase in the post-task stress score (Fig. 4c), as confirmed with individual stress-related variables (Fig. S6f). This is likely explained by an imbalance between fast (freezing) and slow (putative recovery) breathing immobility. Indeed, mice in the SWR disruption group showed a systematic deficit in the duration of slow breathing immobility predicted by the amount of fast breathing immobility (Fig. 4d,e). In turn, this slow breathing immobility deficit predicted the observed increase in stress score (Fig. 4f).

Notably, both groups displayed similar spatial avoidance behaviour, spending comparable amounts of time in the shock zone during conditioning and post-conditioning sessions and receiving the same number of shocks (Fig. 4g-i). The density of shocks received throughout the protocol shows identical dynamics in both groups. Moreover, there was no effect on overall physiology or time spent immobile (Fig. S8h). Thus, SWRs were not required for the learning, expression, or recall of spatial avoidance. These results suggest that SWRs promote slow breathing immobility, and the associated recovery process, contributing to a reduction in post-task stress.

### Recovery state emerges only in safe environments

We hypothesized that the role of SWRs in the slow breathing state is to learn the location of threat-free environments that are suitable for recovery. We therefore examined the time course of breathing frequency throughout all immobility periods in the shock or safe arm. In the example mouse (Fig. 5a), we observed that whereas breathing frequency in the shock arm remained stable throughout the session, the frequency shifted from the fast breathing mode (4–6 Hz) to the slow breathing mode (2–4 Hz) in the safe arm. Comparing breathing frequency during the early and late phases of immobility in each arm showed no change in breathing frequency in the shock arm but a clear drop in the safe arm (Fig. 5b). Similarly, Kendall’s tau, a non-parametric measure of temporal evolution, indicated a decrease exclusively for safe-zone immobility (Fig. 5c). Finally, the linear model predicting breathing rate showed that the contribution of breathing rate emerges gradually throughout the task, consistent with a learning process (Fig. S3c). This change is also observed when quantifying the global physiological variables using the classifier score (Fig. S9a).

**Figure 5.**
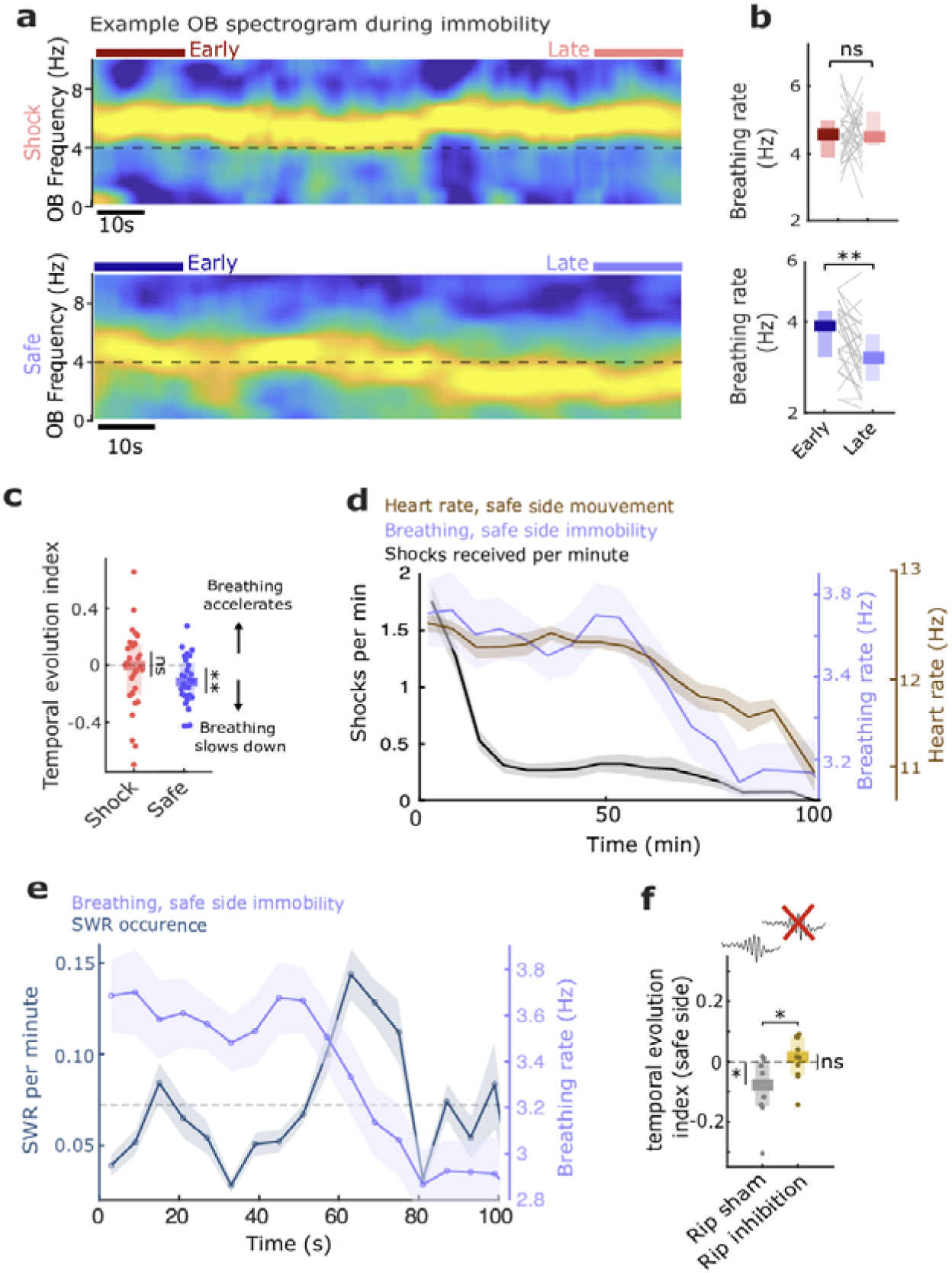
Slow breathing immobility is acquired through safety learning. a. OB spectrum from an example mouse during concatenated data from all shock and safe side immobility throughout conditioning. This shows a stable breathing frequency band on the shock side but a clear reduction of frequency on the safe side. b. Mean breathing frequency during the early and late phases of shock and safe side immobility, showing a difference for safe but not shock arm immobility (n=28/29, **P=0.0042, (NS)P=0.96, Wilcoxon signed-rank test). c. Temporal evolution index of breathing frequency calculated using Kendall’s tau, for shock and safe arm immobility (n=28/29, **P=0.0017, (NS)P=0.63, Wilcoxon signed-rank test). d. Temporal evolution through conditioning of breathing frequency during safe side immobility (blue), heart rate during active periods (orange) and shock number (gray). This shows that avoidance learning, indexed by shock density, occurs faster than safety learning, indexed by reduced heart rate and the appearance of slow breathing frequency (n=29, error bars are s.e.m.). e. Temporal evolution throughout conditioning of the change in breathing frequency during safe side immobility and the number of SWRs. This shows that the drop in breathing frequency co-occurs with an increase in SWR. f. Temporal evolution index for safe arm immobility for SWR disrupted groups and sham groups showing that SWR disruption blocks the evolution of breathing frequency in the safe arm (n=8/8, *P=0.0312, Wilcoxon rank-sum test, individually: *P=0.0312 and (NS)P=0.3750, Wilcoxon signed-rank test). All box plots show median and quartiles. Detailed statistics are provided in supplementary table 1

Heart rate during movement, a proxy for stress during the task (Fig. 2d), decreased throughout conditioning in the safe arm but not the shock arm, independent of speed (Fig. S9b-d). This decrease followed the same dynamics as that of breathing during immobility (Fig. 5d). On the other hand, shock density, which measures avoidance learning, declined significantly faster than the reduction in breathing frequency during immobility and heart rate while moving in the safe zone (Fig. 5d). This suggests that slow breathing immobility only emerges once animals have learnt to distinguish safe from shock zones, as also reflected in their heart rate during locomotion and that this learning process is independent from avoidance learning.

The peak occurrence of SWRs coincided with the decrease of breathing rate during safe side immobility (Fig. 5e). Moreover, SWR disruption prevented the reduction in breathing rate during safe side immobility (Fig. 5f). Thus, the deficit in slow-breathing immobility relative to fast-breathing immobility observed in SWR disruption (Fig. 4d-f) is due to the lack of this transition to slow-breathing immobility in the safe arm.

This suggests the presence of two learning processes with distinct temporal dynamics and underlying mechanisms. The first supports spatial avoidance, likely relying on encoding where the shocks occur, and is independent of SWRs. The second supports the transition to slow breathing immobility in the safe arm, likely relying on encoding where shocks do not occur and that depends on SWRs. These findings suggest that SWRs contribute to safety learning, allowing animals to recognize the absence of threat in the safe zone and implement recovery via slow breathing immobility.

### Diazepam promotes 2-4 Hz recovery state and reduces stress level

If slow breathing immobility supports recovery from stress, then pharmacological agents known to reduce stress could act by promoting this state. To test this, we injected mice with the anxiolytic diazepam (intraperitoneally, 1.5 mg/kg) or a saline control before the U-Maze conditioning (Fig. 6a).

**Figure 6.**
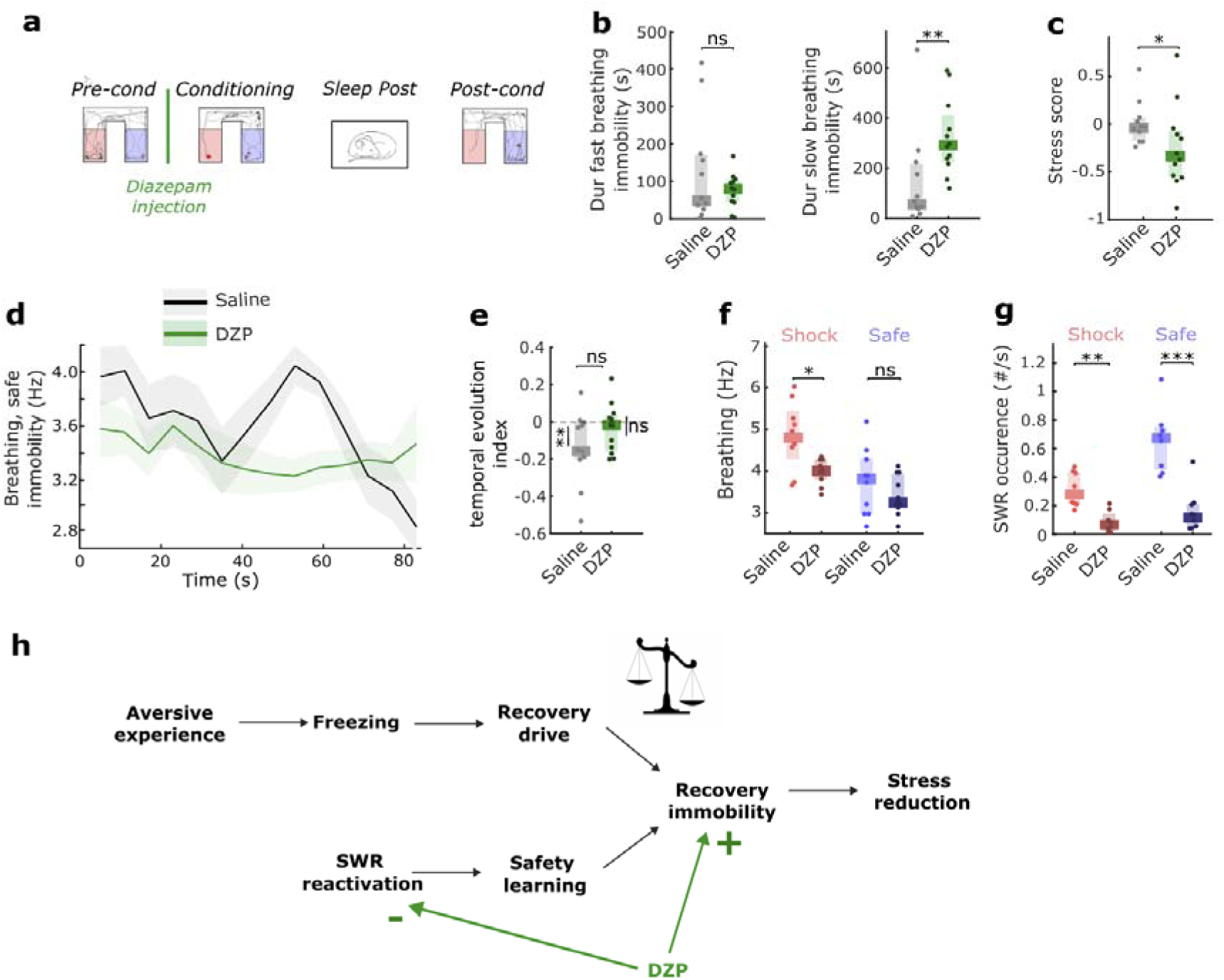
Diazepam increases stress recovery and time in recovery state. a. Scheme showing that diazepam is injected prior to the conditioning session in the U-Maze. b. Diazepam administration led to an increase in the duration of slow breathing immobility but not fast breathing immobility throughout the conditioning session relative to saline controls (n=11/12, (NS)P=0.8294, **P=0.0074, Wilcoxon signed-rank test). c. Diazepam administration led to a reduction in post U-Maze stress relative to saline controls (n=11/12, *P=0.0392, Wilcoxon signed-rank test). d. Temporal evolution through conditioning of breathing frequency during safe side immobility for saline and diazepam injected mice (error bars are s.e.m.). e. Temporal evolution index for safe arm immobility for saline and diazepam injected mice (n=11/12, (NS)P=0.0905, Wilcoxon rank-sum test, individually: *P=0.0098 and (NS)P=0.3394, Wilcoxon signed-rank test). f. Mean breathing frequency during the early and late phases of shock and safe side immobility, showing a reduction in the shock arm for diazepam mice (n=11/12, *P=0.0121, (NS)P=0.5248, Wilcoxon rank-sum test). g. Ripples per second during shock and safe arm immobility show that diazepam systematically reduces their occurrence (n=11/12, **P=0.0019, ***P=5.7589 × 10-4, Wilcoxon rank-sum test). h. Schematic of the proposed homeostatic balance between stress and recovery and the paradoxical action of diazepam which promotes the recovery state but inhibits SWR necessary for safety learning. All box plots show median and quartiles. Detailed statistics are provided in supplementary table 1

After diazepam administration, we observed an overall increase in the duration of slow breathing immobility but no change in the duration of fast breathing immobility (Fig. 6b). Diazepam also led to a reduction in the post-task stress score (Fig. 6c), as confirmed with individual stress-related variables (Fig. S6g). These observations are consistent with an artificially induced “recovery excess” in diazepam mice. Indeed, mice in the diazepam group showed a systematic excess in the duration of slow breathing immobility predicted based on the duration of fast breathing immobility (Fig. S10a,b). In turn, the group-averaged recovery deficit predicted the observed drop in the stress score (Fig. S10c).

This increase in slow breathing immobility in diazepam-treated mice is due to two factors. First, in the safe arm, slow breathing immobility appears immediately without showing any temporal evolution (Fig. 6d,e). Second, during shock arm immobility the average breathing was reduced and shifted toward a physiological state resembling that found in the safe arm for control mice (Fig. 6f, Fig. S10d). Of note, the similar quantities of fast-breathing immobility despite the change in shock arm breathing frequency is due to an overall increase in immobility (Fig. S10e). This shows that after diazepam administration, mice expressed slow-breathing immobility throughout the maze and without learning.

Diazepam did not alter post-conditioning avoidance (Fig. S10f), showing that these effects were not due to spatial memory impairment. Moreover, when diazepam was injected before a control session in the homecage no increase in slow breathing immobility or reduction in stress score was observed (Fig. S10h-j). This rules out a general sedative effect and shows that diazepam-induced changes were specific to aversive conditions.

Interestingly, as previously reported20, diazepam also suppressed SWR occurrence during all behavioural states : immobility (Fig. 6g), homecage quiet wake and sleep (Fig. S10g). Pairing this observation with the immediate appearance of slow breathing immobility in the safe arm, suggests that diazepam bypasses the safety learning necessary for slow immobility to emerge—a process dependent on SWRs in untreated animals (Fig. 5). This aligns with diazepam’s known stress-reducing effects, directly promoting slow-breathing immobility and recovery without requiring the identification of a threat-free area.

## Discussion

Understanding how organisms recover from threat is central to deciphering the biology of resilience. Recovery has previously been operationalized as a passive return to baseline during stress relief periods7 or post-stress sleep8. Our current results identify two distinct types of immobility: a fast-breathing (4–6 Hz) corresponding to classical freezing and a novel low-arousal, slow-breathing recovery state (2–4 Hz). These findings indicate that post-stress recovery is an active process, dependent on a specific behaviour established through hippocampus-dependent safety learning.

This recovery state is characterized by low physiological arousal, with a specific 2–4 Hz slow breathing rate, expressed exclusively in a safe environment after threat exposure, and never observed during baseline, exploratory, or defensive periods. Slow breathing is increasingly recognized for its role in emotional regulation across species. In humans, controlled breathing protocols reduce cortisol levels, enhance parasympathetic tone, and improve stress resilience33,34. In rodents, 2–4 Hz breathing is associated with behavioural calm and the suppression of aversive states12,13,35. Together, these findings strongly support the idea that the slow-breathing state reflects the activation of a specific neural recovery program, rather than general quiescence.

This active recovery state following aversive experience bears a direct resemblance to opponent-process theory. This model of emotional regulation posits that each emotional state is followed by a corrective state of inverse valence36. In our framework, fast-breathing freezing corresponds to the initial defensive process and generates a recovery drive. This drive is alleviated by slow-breathing immobility, a delayed compensatory process that mitigates stress and restores homeostasis.

By showing that hippocampal SWRs play a key role in the expression of slow-breathing immobility, our findings challenge prevailing models of the role of hippocampus in fear that focus on memory encoding. Instead, we reframe this region as a key driver of recovery through safety learning. This learning likely occurs through the reactivation of prior threat episodes observed during SWRs. This would provide a mechanism by which the animal’s model of the environment can be updated to identify non-threatening regions in the absence of further threat exposure. These reactivations offer a potential solution to a key paradox in aversive-based learning: how organisms can learn from the absence of expected punishment.

This novel role for SWRs in safety learning expands their established function in aversive memory consolidation. First, we show that SWRs are not confined to post-learning sleep but instead dominate hippocampal activity during recovery periods within the task itself. Second, previous work has shown that SWRs support the hippocampal-amygdala coordination necessary for encoding fear37, and that their amplification can lead to fear generalization38. Our results demonstrate an opposite role: SWRs enable the identification of non-threatening contexts allowing for fear reduction. Suppressing SWRs, a strategy that has been proposed to reduce traumatic memory consolidation38, may therefore inadvertently impair the capacity for safety learning, increasing vulnerability to stress generalization and relapse. Rather than inhibiting SWRs, interventions that enhance their functional specificity— promoting reactivation in safe contexts—may offer a more effective approach for fostering resilience.

Moreover, by replaying past experiences in a safe, low-arousal context, SWRs may also enable the updating of internal models facilitating cognitive reappraisal. This parallels exposure therapy in humans39, which involves reactivating negative memories in safe contexts—often combined with mindfulness40 or breathing-based techniques33.

Our pharmacological data highlight the importance of this change of perspective on SWR. Administration of diazepam accelerated the onset of the slow-breathing recovery state, bypassing the need for experience-dependent safety inference. However, it also suppressed SWR activity, suggesting a dissociation between behavioural recovery and the safety learning process. This raises key concerns about the long-term efficacy of anxiolytics: while they may offer immediate relief, they could impair the neural mechanisms necessary for enduring emotional regulation.. These results underscore the need for therapeutic strategies that both alleviate acute stress and preserve the endogenous mechanisms of recovery and safety learning.

Finally, our results have significant implications for the interpretation of aversive paradigms and highlight the importance of integrating physiological markers in the study of behavioural neuroscience. First, immobility is used as a canonical measure of fear and uniformly labelled “freezing”41. Using physiological markers, we demonstrate that this immobility includes two physiologically distinct states that are behaviourally indistinguishable: a high-arousal, fast-breathing freezing state (4–6 Hz), and a low-arousal, slow-breathing recovery state (2–4 Hz). Ignoring this distinction may contribute to inconsistencies in interpretation. As a specific example, the literature reports both positive and negative relationships between ‘freezing’—defined as immobility—and circulating corticosterone, used as a marker of stress42–44. If some studies predominantly quantify the high-stress freezing state and others the low-arousal recovery state, then they necessarily will describe opposing relationships with stress hormones. Second, safety learning is studied as the reduction of defensive behaviour (freezing) upon presentation of a stimulus that is never associated with punishment45. Here we identify specific markers for safety: the slow breathing recovery immobility and the reduction in heart rate during locomotion. This provides a new tool to investigate the specific behavioural manifestations of safety learning and its functional importance in stress reduction.

By identifying a distinct low-arousal recovery state associated with SWR-mediated reactivation of aversive memories, our study offers a novel framework for understanding resilience. These findings suggest that recovery is not simply the absence of fear, but rather a positively defined neural process with specific physiological and neural correlates. Importantly, they redefine the role of hippocampal replay in aversive learning—not only as a mechanism of memory consolidation, but as a substrate for emotional regulation with broad implications for our understanding of stress-related disorders and their treatment.

## Supporting information

Stats supplementary

## Acknowledgments

The authors thank members of the laboratory for their valuable feedback on the manuscript. We gratefully acknowledge financial support from the ERC Consolidator Grant, the Agence Nationale de la Recherche (ANR), ESPCI Paris – PSL, and the CNRS.

## Author contributions

B.M., S.B., and K.B. conceived and designed the experiments and analyses, analyzed the data, and wrote the manuscript. D.B., C.H., M.C., E.C., and C.S. performed the experiments. T.G. and K.B. supervised this work.

## METHODS

### ANIMALS AND SURGICAL PROCEDURES

#### Animals

All behavioural experiments conformed to the official European guidelines for the care and use of laboratory animals (86/609/EEC) and the policies of the French Committee of Ethics (Decrees n° 87-848 and n° 2001-464). The animal housing facility at the laboratory where the experiments were conducted is fully accredited by the French Direction of Veterinary Services (B-75-05-24, 18 May 2010). C57Bl6/J male mice (approximately 27-32 g, 3-6 months of age; The Jackson Laboratory) were used in electrophysiological recordings. Mice were housed in an animal facility with a light cycle from 08:00 to 20:00, with *ad libitum* access to food and water. Temperature and humidity in the room were kept at 20-24°C and 40-60%, respectively. Mice were housed with a maximum of four per cage before surgery and individually afterward.

#### Surgical protocols

Premedication consists of subcutaneous injection of buprenorphine at 0.1mg/kg and a local injection of lidocaine at 5mg/kg. Mice were then anaesthetized with isoflurane (1-3%) and placed on a heating pad during the entire surgery and Ocry-gel® was regularly applied to avoid ocular drying. Stereotaxic coordinates were taken following stereotaxic guidance after which craniotomies were performed. Mice were implanted with recording electrodes (tungsten wires with PFA isolation) in the right olfactory bulb (coordinates: AP +4, ML +0.5, DV −1.5) and the right CA1 hippocampal layer (coordinates: AP -2.2, ML +2.0, DV −1.0). LFP were recorded using. To record the electrocardiogram, a highly flexible coiled wire (small gauge biopotential leads, DSI) was sutured to the thoracic muscles above the heart and then travelled under the skin to be fixed to the head-stage and preamplified with the other recording wires. In a subset of mice, we also recorded a nuchal electromyogram by suturing silver wires in the muscles behind the skull. In order to provide aversive stimulation via an eyelid shock, loops of exposed silver wire were sewn through the tissue anterior or posterior to each eye, ensuring no wire was accessible after suturing the skin. We also used dPAG aversive stimulation with bipolar electrodes implanted in the dPAG (60-mm-diameter stainless steel, AP -3.52, ML 0.5, DV -2.7). The mice in the ripples inhibition protocol were also implanted, bilateral bipolar electrodes were implanted in the ventral hippocampal commissure (coordinates: AP -0.8, ML ±0.5, DV -1.9). On a subset of mice, silicon probes (ASSY-236 E-, 64ch) were implanted above the

CA1 pyramidal layer of the right hippocampus (AP +2.2, ML +2.0, DV −1.0) to record single units. The probes were progressively lowered until they reached the CA1 pyramidal layer, where ripples were recorded. Probes, wires and the EIB were secured to the skull using dental cement (Superbond Universal). At the end of surgery, mice received a subcutaneous injection of meloxicam (10 mg/kg), glucose and NaCl and placed on a heating pad until recovering locomotion. After surgery, mice were monitored daily to verify recovery and recordings began 1 week after surgery, when animals had recovered normal behaviour.

### BEHAVIOURAL PROTOCOLS

#### Recording setup

Signals from all electrodes were recorded using an Intan Technologies amplifier chip (RHD2216, sampling rate 20 kHz). The Intan RHD2132 on the animal’s head is equipped with a 3-axis accelerometer, offering high temporal resolution data on head movements. LFP were stored at 1250 Hz. Animals were tracked using an overhead thermal camera (FLIR A325sc, Teledyne FLIR LLC, Oregon, USA) and custom MATLAB-based software (The Mathworks, Inc, Massachusetts, USA), which identified the center of mass of the hottest spot in the camera field of view as a mouse. Acquisition rate of the camera was 15 Hz. Electrophysiology and tracking data were synchronized using Arduino-based code, which managed TTL pulses to log start and end of each recording session.

#### Aversive stimulation calibration

The eyelid shock was delivered between the anterior and posterior wires on each eye so that the mouse received a simultaneous eyelid shock on both sides. It was composed of 1ms bipolar pulses, delivered at 140 Hz for 100 ms. The dPAG stimulation consisted of pulses with identical features delivered over a 200 ms duration. Stimulation was delivered using a PulsePal stimulator (Sanworks). To calibrate the stimulation voltage of aversive stimulations, mice were submitted to sessions of 130 s where they received 4 shocks. Voltage was set to 0V for the first session and was increased to 1V between each session. The first amplitude to evoke both jumping and freezing was used for the conditioning. In general stimulation between 2 and 4V evoked reliable behavioural responses. These aversive calibrations were realized the day before the U-Maze experimental day, in a squared box of 20x20cm.

#### U-Maze environment

The U-Maze consists of arms measuring 19 cm in width and 26 cm in length, with a center zone forming a rectangle of 12 cm in width and 46 cm in length. The walls of the maze provide visual cues: one side is stripped whereas the other has a zebra motif and large colored shapes at the end of each arm. Between each session walls and floor are cleaned either with 30% ethanol or laboratory-purpose detergent (ANIOS spray). The shock and safe arms could be closed off with transparent doors that could be slid in and out of the maze quickly in order to block the animal in these arms in a subset of sessions.

#### U-Maze conditioning protocol

Before beginning the protocol for each mouse, one arm is chosen at random to be the shock or safe arm. At the beginning of each session in the U-Maze, the mouse is placed in the center arm, facing towards the shock or safe arm in alternating order. The mouse is first habituated to the environment during 4×15 min sessions without any shocks (*Habituation*). During the habituation sessions, transparent doors are added / removed from the maze while the animal is in different positions to habituate to the different maze configurations. The mouse is then placed in its homecage where it stays until it has slept for a total of one hour (*Sleep Pre*). The mouse is then placed in the environment for 4x3 minutes sessions which are used to quantify baseline exploratory of the arms (*Pre-conditioning).* The conditioning then begins, during which the mouse’s position is detected automatically by custom Matlab software which delivers an aversive stimulation 13s ±1s after the mouse enters the shock zone. If the animal remains in the zone the shock is repeated with the same temporal delay. During each 8min conditioning sessions, mice are either free to navigate throughout the whole maze (*Conditioning Free)* or are blocked by a transparent door in the shock or safe area for the first 5 minutes when the door is removed leaving the mouse free to explore during the last 3 minutes (*Conditioning blocked shock / Conditioning blocked safe).* During the *Conditioning blocked shock* sessions, the mouse receives 4 shocks while blocked at fixed times (120/150/180/210 s). After conditioning, the mouse is then placed in its homecage where it stays until it has slept for a total of one hour (*Sleep Post*). The mouse is then placed in the environment for 4x3 minutes sessions which are used to quantify post-conditioning occupation of the arms (*Post-Test).* The mouse is then blocked in the shock arm and receives a single shock (*Imposed shock)*. It is then blocked alternatively in the shock or safe arm during 2 sessions for each arm, this allows measuring extra immobility time (*Recall shock / Recall safe)* (Fig. S2a).

Based on this protocol structure, the number of conditioning sessions and the use of transparent doors was adapted to the different experimental requirements, the mice submitted to each protocol are listed in table 1:

- *Blocked sessions, long* : This protocol allows to characterize extensively the two types of immobility and in particular their temporal evolution. First, we performed a total of 12 sessions. We performed 6 before and 6 after a sleep period, in order to ensure the animal wasn’t tired during the protocol. Second, we equalized the time spent in shock and side immobility by introducing sessions in which mice are blocked in each arm by a transparent wall to which they had been previously habituated. This compensates for the low shock side immobility when free to escape observed in eyelid shock mice (Fig. S2b,c)
- *Blocked sessions, short* : This protocol is similar to the long protocol but we only performed 9 conditioning sessions without an intervening sleep session. This reduced the time required to complete the protocol for the experimentally demanding ripple inhibition protocol.
- *No blocked sessions, short* : This protocol is adapted for mice with dPAG stimulation who show shock and safe side immobility with the same characteristics as eyelid shock mice (Fig. S2a) but display long duration of immobility without being blocked in either arm (Fig. S2b,c). Thus, performing 5 conditioning sessions without ever blocking the animal yielded similar total time immobile for the dPAG animals as those with the long eyelid shock protocol (Fig. S2c). The advantage of this protocol is that we could record in a short period of time large amounts of neural data during freezing which guaranteed spike sorting stability throughout the session.

**Table 1:** Mice groups.

| Experiment | Figure | Aversive stimulation | Conditioning protocol | Size |
| --- | --- | --- | --- | --- |
| U-Maze characterization | Fig. 1-2-3-5 except from SU | Eyelid shocks | U-Maze (blocked sessions, long and short) | n=29 |
| U-Maze characterization - alternative stimulation (dPAG) | Fig. S2 | dPAG | U-Maze (no blocked sessions, short) | n=20 |
| Hippocampal reactivation study | Fig. 3 SU only | dPAG | U-Maze (no blocked sessions, short) | n=9 |
| Ripple inhibition | Fig. 4 | Eyelid shocks | U-Maze (blocked sessions, short) | n =10 inhibition<br>n=10 control |
| Diazepam administration | Fig. 6 | Eyelid shocks | U-Maze (blocked sessions, long) | n=13 diazepam<br>n=15 saline |
| Tone fear conditioning | Fig. S4 | Electrical footshock | Tone conditioning | n=14 |
| Context fear conditioning | Fig. S4 | Electrical footshock | Contextual conditioning | n=5 |
| Social defeat stress | Fig. 2 and supp Fig. S2 | Encounter with dominant male | Social defeat stress | n=13<br>n=14 |

#### Tone fear conditioning

On day 1, mice were habituated to the testing environment without sounds. On day 2, mice were submitted to a habituation session in the conditioning context during which the CS− and the CS+ were presented 4 times. Discriminative fear conditioning was performed on the same day by pairing the CS+ with a US (1-s foot-shock, 0.6 mA, 8 CS+ US pairings; inter-trial intervals, 20-180 s). The onset of the US coincided with the offset of the CS+. The CS-was presented after each CS+ US association but was never reinforced (8 CS− presentations; inter-trial intervals, 20-180 s). On day 3, conditioned mice were sub-mitted to a test session during which they received 4 and 12 presentations of the CS− and CS+, respectively. The conditioned sounds consisted of a 30 s sequence of 50 ms pips repeated 27 times at 80 dB sound pressure level. The pips of the CS− and CS+ could be either 7.5 kHz or white-noise and were counterbalanced across experimental groups. Different environments were used for conditioning and testing to avoid any contributions of contextual conditioning, they differed in shape (cylinder vs square), colour (black vs white), floor (metal grid vs plastic) and odour cues (70% ethanol vs 1% acetic acid). Fear conditioning experiments were performed using the Imetronic fear conditioning system.

#### Contextual fear conditioning

On day 1, mice were habituated to control environment A and B for 30 minutes. On day 2, mice were submitted to a conditioning session of 30 minutes in environment C during which they received electrical foot shocks (1 s, 0.6 mA, 8 CS+ US pairings; inter-trial intervals, 20-180 s). On day 3, conditioned mice were put back in environment A, B and C for 6min per environment.

#### Calibration of SWRs detection and disruption

vHC stimulations were composed of a biphasic pulse, with each phase lasting 0.2 ms and an inter-pulse interval of 0.1 ms with a PulsePal stimulator (Sanworks). To detect SWR occurrence in real time, electrophysiological recordings from the hippocampal CA1 pyramidal layer were converted to a digital-to-analog channel (DAC). This signal was band-pass filtered (120-250 Hz) using an analog filter (Alligatortech®) and then transmitted to a pre-amplifier and data acquisition interface (CED 1401). A Spike2 (v6) software interface was utilized to manually set a customized threshold for each mouse on the band-pass filtered signal. In order to ensure that the SWRs was not contaminated by muscular artefacts that can occur within the same frequency band, a threshold was also set on a non-pyramidal CA1 channel without SWRs. When the threshold on the SWRs channel but not the non-SWRs channel is exceeded, a TTL was sent via Spike2 sequencer to a PulsePal stimulator (Sanworks). Overall, the median duration of stimulation delay was 12 ms. In order to calibrate vHC stimulation voltage, a calibration session was performed the day before mice explored the U-Maze when mice were in NREM sleep. During these sleep sessions, we initially recorded without stimulation to characterize the baseline SWRs features. Subsequently, stimulation at 0V with a 150-250 ms delay following SWRs detection was applied. This step ensured that vHC stimulations were properly triggered upon SWRs detection without altering the waveform massively, serving as a control for accurate stimulation timing. Following this validation phase, we conducted a series of sessions where vHC stimulations were delivered immediately upon SWRs detection, with increasing voltage levels. Each session lasted 130 seconds, with the stimulation voltage incremented by 0.5V between sessions. The first voltage that produced SWR disruption and a clear evoked potential was selected for subsequent experiments.

#### SWR disruption during U-Maze conditioning

During all conditioning sessions, vHC stimulation was initiated either immediately after SWRs detection (Ripple inhibition group) or with a random delay ranging from 180-220 ms after detection (Ripple control group). This stimulation was gated by the animal being immobile based on real-time camera-based tracking. Mice that exhibited ictal or epileptic-like events during either calibration or U-Maze experiments were excluded from the analyses. These events were rare, occurring in only 2 out of the 22 mice prepared for this study.

#### Social defeat stress

The experiment started at 9am. First, the intruder (experimental mouse) was introduced into the home cage of an unfamiliar, aggressive male CD1 resident mouse for 5 minutes for physical interactions. If the conflicts were too intense, the mice were briefly separated for a few seconds to avoid wounds and damages to the implant. In a second step, the two mice were separated by a transparent and perforated partition placed in the middle of the CD1 mouse home cage for 20 minutes. During this step, the experimental mouse remained in sensory contact (olfactory, visual and auditory) with the CD1 mouse. In a final step, a second sensory exposure of 20 minutes was repeated in the experimental mouse’s home cage to evaluate the effects of exposure in a safer environment. During the stress procedure the experimental mouse usually shows clear submissive posture and freezing behaviour. The CD1 mouse was then removed from the cage and the experimental mouse. The experimental mouse was then connected to the recording cable and allowed to sleep for the rest of the day (in its home cage with the partition). The sleep sessions usually started around 10 am up to 6:30 pm.

#### Drug delivery

Diazepam (Sigma-Aldrich®) was prepared as a suspension in physiological saline with 0.1% Tween 80 and administered intraperitoneally prior to conditioning sessions. The anxiolytic efficacy of the dose was confirmed by an increase in open arm time in the elevated plus maze.

### DATA ANALYSIS

#### General code and software

Analyses were performed with custom made Matlab programs, based on generic code that can be downloaded at www.battaglia.nl/computing/ and http://fmatoolbox.sourceforge.net/. Recordings were visualized and processed using NeuroScope and NDManager (http://neurosuite.sourceforge.net/).

#### Behavioural analysis

The instantaneous position of the mouse was extracted from the overhead thermal camera frames by thresholding on the temperature (since the mouse is markedly hotter than its environment). Due to the fact that recordings were made in different rooms in slightly different visual settings, all raw trajectories were aligned to common coordinates based on the found corners of the maze. To assess freezing behaviour, we used the instantaneous variation in the 3-axis accelerometer measure which provides high temporal resolution data on head movements. Freezing was defined as episodes lasting at least 2 seconds, during which the accelerometer readings stayed below a predefined threshold uniformly set for all mice. For mice without an accelerometer on the headstage, freezing behaviour was determined using a quantity of movement, defined as the pixel-wise difference between two consecutive frames. In order to verify that now periods of sleep or strong drowsiness were included, epochs of immobility with OB gamma power below the wake/sleep threshold (as determined by sleep scoring on a previous homecage sleep session) were excluded from the analysis since this parameter has been shown to reliably differentiate wake immobility from sleep^23^. These excluded epochs constituted only a minor portion of the data, mainly occurring at the end of the protocol during recall sessions at the end of the day. Throughout the paper we quantify behaviour in the shock zone (the shock arm up to the corner, corresponding to the zone in which mice could receive shock) and the safe zone (the safe arm and the closest corner since some mice were mainly immobile in this region, with immobility features identical to those in the safe arm).

#### LFP analysis

Electrophysiological data were analyzed using custom-written MATLAB scripts (The MathWorks Inc., Natick, MA, 2000) incorporating the Chronux toolbox (http://www.chronux.org). Spectrograms were generated using multi-taper Fourier analysis to minimize finite windowing effects. For all analyses, five tapers and a window size of 3 seconds with a 0.2-second shift between bins were applied. For each mouse, the spectrograms of all freezing episodes were concatenated and then averaged to obtain a representative spectral profile. The mean spectrum was computed as the average power at each frequency and subsequently normalized for each mouse using the peak power during shock freezing as the reference value.

#### Breathing rate extraction

It is well documented that the olfactory bulb LFP tightly tracks breathing^46^. Instantaneous breathing was calculated from OB LFP spectrograms, which were binned every 0.2 seconds. Within each bin, the frequency between 1-8 Hz corresponding to the maximum power was identified as the breathing frequency.

#### Heart beat analysis

Individual heartbeats were detected through a two-step process. First, the signal was band-pass filtered to isolate the relevant frequency range, and large heartbeats were identified as events peaking above 3 standard deviations (s.d.) from the mean. These large events were used to create a template, which was then convolved with the original signal. In the second step, all heartbeats were detected by applying a threshold of 1.5 s.d. to this convolved signal. This approach allowed for the detection of smaller events that matched the correct shape of heartbeats while avoiding large events that did not conform to the expected shape. Heart rate corresponds to the inverse of the inter-beat time. Heart rate variability was calculated as the standard deviation of a sliding window of five heartbeat measures. When comparing heart rate between different mice, we used a value normalized to a baseline period of calm in a stress-free environment (the homecage) because of the substantial variability between individuals.

#### Electromyography

EMG data was filtered in the 50-300 Hz band and instantaneous amplitude derived from the Hilbert transform. This time series was then smoothed using a 2 s sliding window.

#### Tail temperature extraction

Mice were tracked from above using an infrared camera (A325 FLIR). To retrieve tail temperature, we used DeepLabCut^TM^ with markers at the base and the middle of the tail. We measured along the perpendicular axis passing by the middle of these two markers the temperature maximum that was defined as tail temperature.

#### Offline SWRs detection

Ripples detection was performed by band-pass filtering (120-250 Hz), squaring and normalizing the field potential recorded in the CA1 pyramidal layer. Two methods were used to detect SWRs. The first one identified SWRs as periods of time where filtered signal exceeded 4 standard deviations (s.d.) with the peak value exceeding 6 s.d. The second one labeled periods of time where squared filtered signals exceeded 2 s.d. with the peak value exceeding 5 s.d. For both methods, STD was calculated on the whole day recording. For both methods, minimal duration of SWRs was set to 20 ms, maximal duration was set to 200 ms, and minimal inter-ripple interval was 15 ms. Results of both methods were merged to obtain the final set of SWRs. Ripples were defined as events peaking above 6 s.d. The beginning and end of a SWR were defined with a threshold of 3 s.d. Events shorter than 20 ms and longer than 200 ms were removed.

#### States-scoring

We employed the sleep scoring method described in^23^. The primary difference from other conventional sleep scoring methods is the use of OB gamma oscillations to determine sleep or wake states instead of EMG or movement quantity. OB gamma power values (50-70 Hz) were fitted to a mixture of two Gaussian distributions. A manual threshold was applied, with values below the threshold classified as sleep and those above as wake. Periods of sleep and wake shorter than 3 seconds were merged into adjacent periods to avoid artificially short epochs. REM and NREM sleep distinctions were made using the theta/delta (5-10 Hz/2-5 Hz) ratio of the hippocampal signal, similar to other scoring methods. Quiet wake was defined using the same criteria as for freezing in the home cage, based on headstage acceleration during the epoch preceding the first sleep bout lasting more than 30 seconds.

#### Stress score

We measured post-task stress levels using four indicators previously described in the literature : percent time awake (since stress induces insomnia)^26^, percent of sleep time in REM (since stress reduces REM sleep)^26^, thigmotaxis quantified as the percent of time near the walls (since stress increases thigmotaxis)^27^ and heart rate (since stress increases heart rate)^28^. These four parameters are strongly correlated and therefore we combined them into a single stress score using PCA. We performed PCA on the 4 parameters measured for all mice and then projected the data onto the first eigenvector which explained 60% of variance and has positive loadings for parameters that increase with stress and negative loadings for parameters that decrease with stress. We used this value as the “stress score”. When one of the parameters had not been recorded for a mouse, the stress score was calculated using the others.

#### Slow breathing deviation

As explained in the main text, we propose that fast breathing immobility corresponds to classical freezing which is expected to enhance the animals stress level as it requires a strong metabolic and cognitive adaptation to the current level of high threat and that slow breathing immobility corresponds to a state of recuperation. It follows from this that animals showing more fast breathing immobility relative to slow breathing immobility are expected to have an excess of fear-related stress and a recovery deficit and conversely animals showing less fast breathing immobility relative to slow breathing immobility are expected to have a deficit of fear-related stress and a recovery excess. We quantified this as a “recovery deviation” by fitting a linear function the relation between the amount of fast and slow breathing immobility and then quantifying for each animal the difference between the measured duration of slow breathing immobility and that predicted by the linear model.

#### Linear classifier of immobility states

To provide a global view of the difference in physiology between shock and safe side immobility, we combined four parameters (breathing frequency, heart rate, heart rate variability, OB gamma power and frequency). We trained linear classifiers using leave-one-out cross-validation (train on n-1 mice, test on mouse n) to classify shock and safe side immobility based on these parameters. This provides both a measure of overall accuracy (what % of mice had correctly classified shock and safe side immobility) and a score for each immobility type that measures its distance from the hyperplane best separating the two immobility classes. The more positive this score the more the classifier is confident that the immobility is “shock-like” and the more negative the more “safe-like”.

#### Temporal evolution index

To assess the temporal evolution of the respiratory rate, a Kendall’s tau (τ) correlation coefficient was computed using spectrogram values. For each mouse, the respiratory rate was previously estimated from the spectrogram data, binned every 0.2 s, resulting in a time series of respiratory rate values. Kendall’s tau is a non-parametric measure of the monotonic relationship between two variables, in this case, time and respiratory rate. Mathematically, τ is calculated as:

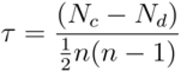

where Nc is the number of concordant pairs, Nd is the number of discordant pairs, and n is the total number of observations. Pairs of observations (x_i_,y_i_ and x_j_,y_j_ with i < j) are concordant if y_i_>y_j_ when x_i_>x_j_, and discordant otherwise. A τ value of +1 indicates a perfect increasing monotonic trend, -1 indicates a perfect decreasing trend, and 0 suggests no monotonic relationship. Statistical significance was set at p < 0.05. This approach quantitatively identified temporal patterns in the respiratory rate, including increases, decreases, or stability throughout the experimental session.

#### ROC analysis

To quantify the relationship between a physiological parameter value and the type of freezing, we used Receiver Operator Characteristic (ROC) curves (Dayan & Abbott, 2014). This analysis assesses the ability of an ideal observer to predict whether an animal’s freezing is based purely on the physiological parameter value. Each point on the ROC curve represents the true positive rate and false positive rate for different threshold values z over 2-second bins of freezing. The ROC curve and its area under the curve (AUC) measure the procedure’s performance, indicating the probability that an ideal observer can discriminate between shock-induced and safe-induced freezing during a 2-second period. An AUC of 0.5 means the physiological parameter provides no information about the freezing type, while an AUC of 1 indicates perfect predictability.

#### Spike sorting

For spike sorting, extracted waveforms were sorted via a semi-automated cluster cutting procedure using KlustaKwik and Klusters (http://neurosuite.sourceforge. net/). We then performed manual verification of single-unit quality based on waveform, firing stability and refractory period.

#### Place cell identification

Rate maps for all recorded single units were generated with a spatial bin of 0.8 cm during epochs when the animal’s speed exceeded 3 cm/s. These maps were then smoothed using a 2D Gaussian kernel with a standard deviation of 2. A single unit was classified as a place cell if its spatial information was greater than 1 bits/s during moving epochs of the considered session (where speed was greater than 2 cm/s). Bin size was 5 ms. These place cells were classified into shock, safe and centre preferring cells by identifying the location of their firing peak in the U-Maze.

#### Measure of reactivation strength

To assess the similarity between patterns of population-level activity, we utilized the “reactivation strength” measure developed by Peyrache et al.^47^. Neural firing activity is initially sampled from a template period and then decomposed using PCA into components that each capture a specific pattern of neural activity based on the neuronal correlation matrix. These identified patterns are then used to scan a match period to detect times when population activity resembles that of the template period, by calculating the instantaneous reactivation strength for each pattern. For instance, in our study, ripple-evoked activity (“during ripples”, ±50 ms around ripples peak detection) served as the template epoch, and reactivation strength was calculated between the averaged activity of spikes of the corresponding epoch, here particularly moving epoch when speed assessed by camera was considered above 2 cm/s. Mean similarity score corresponds to the reactivation strength between the template epoch and the activity of hippocampal A control template (“outside ripples”) corresponds to the time +1.9-2s after ripples peak detection. Bin size was 100 ms.

Let Q be the z-scored (n x b) matrix of firing rates for the n recorded units over b time bins during the match period. The correlation matrix C is defined as:

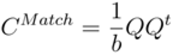

The matrix C can be expressed using eigen decomposition as a sum of projectors P, weighted by their associated eigenvalue λ:

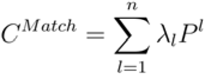

Each projector P captures a specific pattern of correlated activity among the population of neurons, and λ indicates how much of C is captured in λ. Only projectors P associated with eigenvalues λ larger than λmax, as defined in Peyrache et al., (2010), are retained as significant patterns of activity. These projectors can then be used to search the match period for peaks in reactivation strength for the associated pattern:

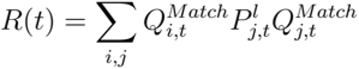

In all figures, the mean reactivation strength is shown, averaged over all significant templates.

#### Linear model of breathing frequency

A linear model was constructed to predict the breathing frequency during immobility in the conditioning phase based on various predictors in order to test their contribution. One model was fitted for each mouse. The predictors we tested were :

- total time spent immobile
- linearized position in the maze (0 to 1 from shock to safe arm)
- global time in conditioning
- time since last shock
- mouvement prior to immobility

In some cases, prior analysis revealed that these predictors had a clear non-linear relation to breathing frequency and so were transformed prior to fitting.

First, the time since the last shock (t_s_) was applied to a negative exponential (Fig. S3d), according to the formula :

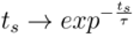

The parameter D was optimized using a grid search with five-fold cross-validation to select the set of hyperparameters that maximized the coefficient of determination of the model.

Second, position (x) was transformed using a sigmoid, in order to account for the non linearity mapping of the breathing frequencies along the U-Maze :

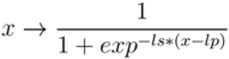

The parameters ls=20 and lp=0.5 were fixed, lp imposes where the sigmoid shifts from 0 to 1 and ls the steepness of this transitions

Finally, given that early in conditioning safe and shock side breathing were similar and safe arm immobility showed a clear temporal evolution we also included a position predictor that took this into account by multiplying the position predictor by a sigmoid function of global time (t_g_). Intuitively, this means that early in conditioning the sigmoid of global time is 0 so position plays no role in breathing frequency and then this variable transitions to 1 and position impacts the animals breathing frequency. In this variable, time was transformed using a sigmoid :

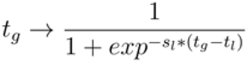

The parameter t_l_ can be seen as the time of learning and s_l_ as the steepness of the learning transitions, both were optimized using a grid search

Mice with less than 100 data points or for which we could not achieve a positive coefficient of determination after optimisation of the parameters were excluded from the analysis.

To determine the contribution of individual predictors on the variability in breathing frequency during immobility, we calculated the prediction gain for the targeted variable in the linear model. This gain was defined as:

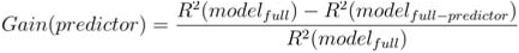

In this formula, the full model is a linear model that includes the following predictors: total time spent immobile, sig(position) **×** sig(time in conditioning), exp(time since last shock), time in conditioning, and movement quantity. The model “full - predictor” is a model fitted while omitting the given predictor, meaning that the coefficients of the model are fitted again without the given predictor. Nevertheless, the hyperparameters for the transformations extracted from grid search are kept while fitting omitted models. This formula therefore enables the evaluation of each single predictor’s contribution in explaining the variance in the breathing frequency.

#### Statistics

For each statistical analysis in the manuscript, the Kolmogorov-Smirnov normality test was first conducted on the data. If the data did not meet the normality criterion, non-parametric tests were utilized. Consequently, we represent the median and quartiles of the data in boxplots in all figures, aligning with the use of non-parametric tests. For comparing proportions, the standard chi-squared test was employed. A two-way mixed repeated ANOVA was conducted using GraphPad Prism software (GraphPad Software, La Jolla, California, USA). For Wilcoxon-Mann-Whitney test and Wilcoxon signed-rank test, we report the signed rank statistic when the number of replicates is insufficient to provide the normal Z statistic. Pearson correlation was used for samples ≥30. For sizes 10–29, Shapiro-Wilk tests determined normality; Pearson was used if normality held, otherwise Spearman. Spearman was applied for samples <10. Correlations were considered significant if their p-values were less than 0.05. For repeated correlations across a variable such as time, Bonferroni correction was applied with α=0.05. The trend line was determined using the least-squares method.

## Supplementary materials

**Figure S1.**
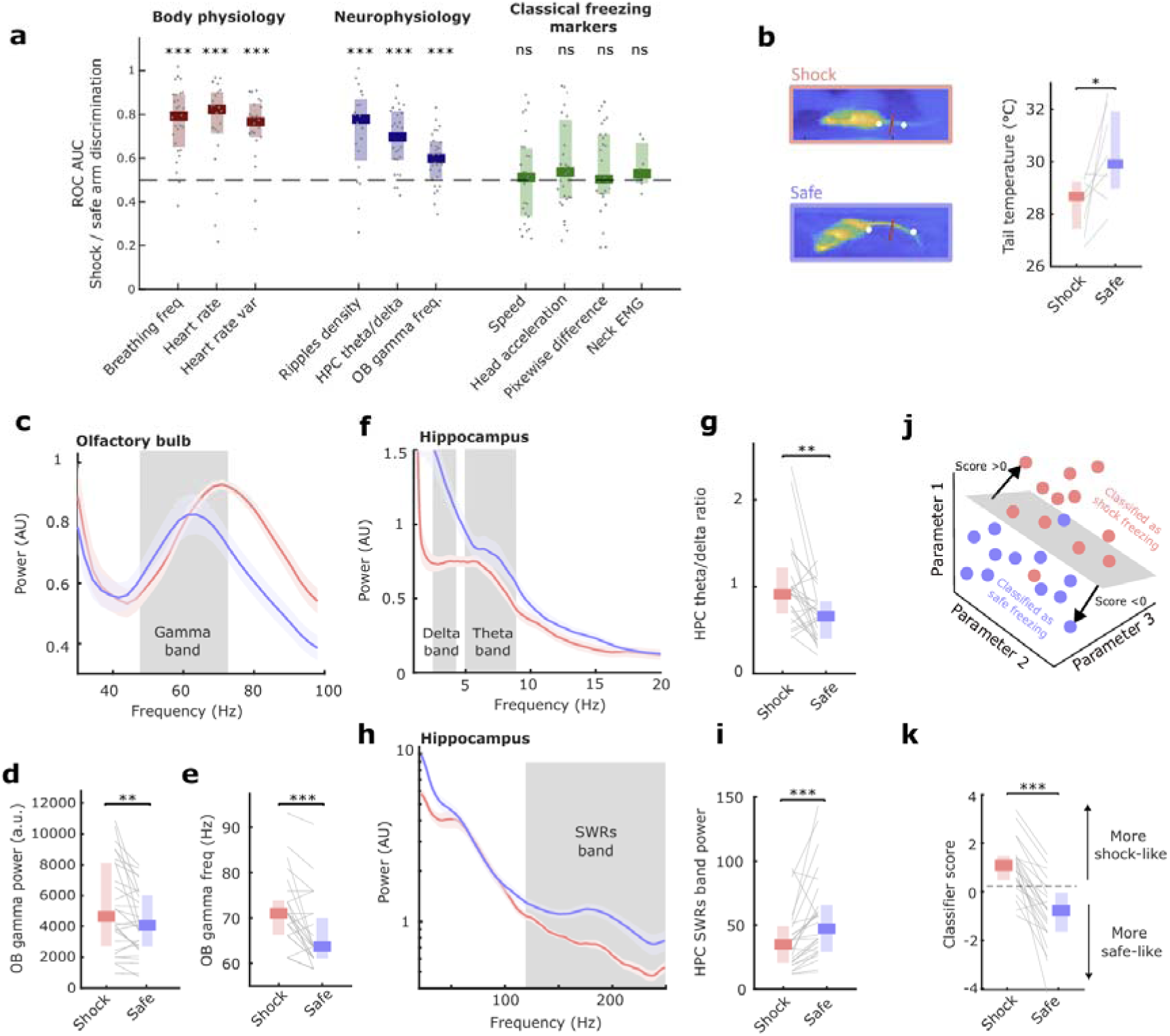
Details of physiological characterization of two immobility states. a. Area under receiver operating curve (ROC) that quantifies the ability of each variable to discriminate shock and safe arm immobility. This shows that markers of body and brain physiology discriminate between the two well (>0.5) but that traditional markers used to identify freezing do not. Both states would therefore likely be classified as ‘freezing’ in previous studies (n=28, 22, 22, 22, 20, 27, 28, 27, 28, 27, 7, ***P=7.2 **×** 10^−6^, 2.59 **×** 10^−4^, 1.76**×** 10^−4^, 8.75**×** 10^−4^, 8.91**×** 10^−4^, 2.91**×** 10^−5^, 6.91**×**10^−4^, (NS)P=0.91, 0.24, 0.17, 0.21, Wilcoxon signed rank test). b. Thermal image showing measurement of mouse tail temperature (left) and mean tail temperature during shock and safe side immobility (right). This shows that temperature is reduced on the shock side, which is indicative of blood flow directed towards hindlimbs in preparation for flight (n=8, *P=0.0391, Wilcoxon signed rank test). c. Olfactory bulb spectrum during shock and safe side immobility, error bars are sem. d-e. OB gamma (50-70 Hz) power (d) and peak frequency (e) during shock and safe side immobility, showing reduced power and frequency on the safe side which may be indicative of lower arousal (n=28, *P=0.0391, Wilcoxon signed rank test). f. Hippocampus low-frequency spectrum during shock and safe side immobility, showing slow theta peak on the shock side. Error bars are sem. g. Hippocampal theta (5-8 Hz) to delta (2-4 Hz) band power ratio during shock and safe side immobility (n=27, ***P=2.6818 **×** 10^−5^, Wilcoxon signed rank test). h. Hippocampus high-frequency spectrum during shock and safe side immobility, showing increase in ripple band power in the safe side. Error bars are sem. i. Hippocampal sharp wave ripple band (120-250 Hz) power during shock and safe side immobility (n=26, ***P=6.3294 **×** 10^−5^, Wilcoxon signed rank test). j. Schematic showing how a linear classifier separates periods of immobility in the shock and safe side arm using multiple parameters. k. Classifier scores for immobility on the shock and safe sides across all mice. The classifier was trained using a leave-one-out cross-validation approach: for each mouse (n), the model was trained on the data from the remaining (n–1) mice, and the resulting classifier was used to predict the immobility score for the held-out mouse on both the shock and safe sides. This procedure was repeated for every mouse (n=22, ***P=4.6125 **×** 10^−5^, Wilcoxon signed rank test).

**Figure S2.**
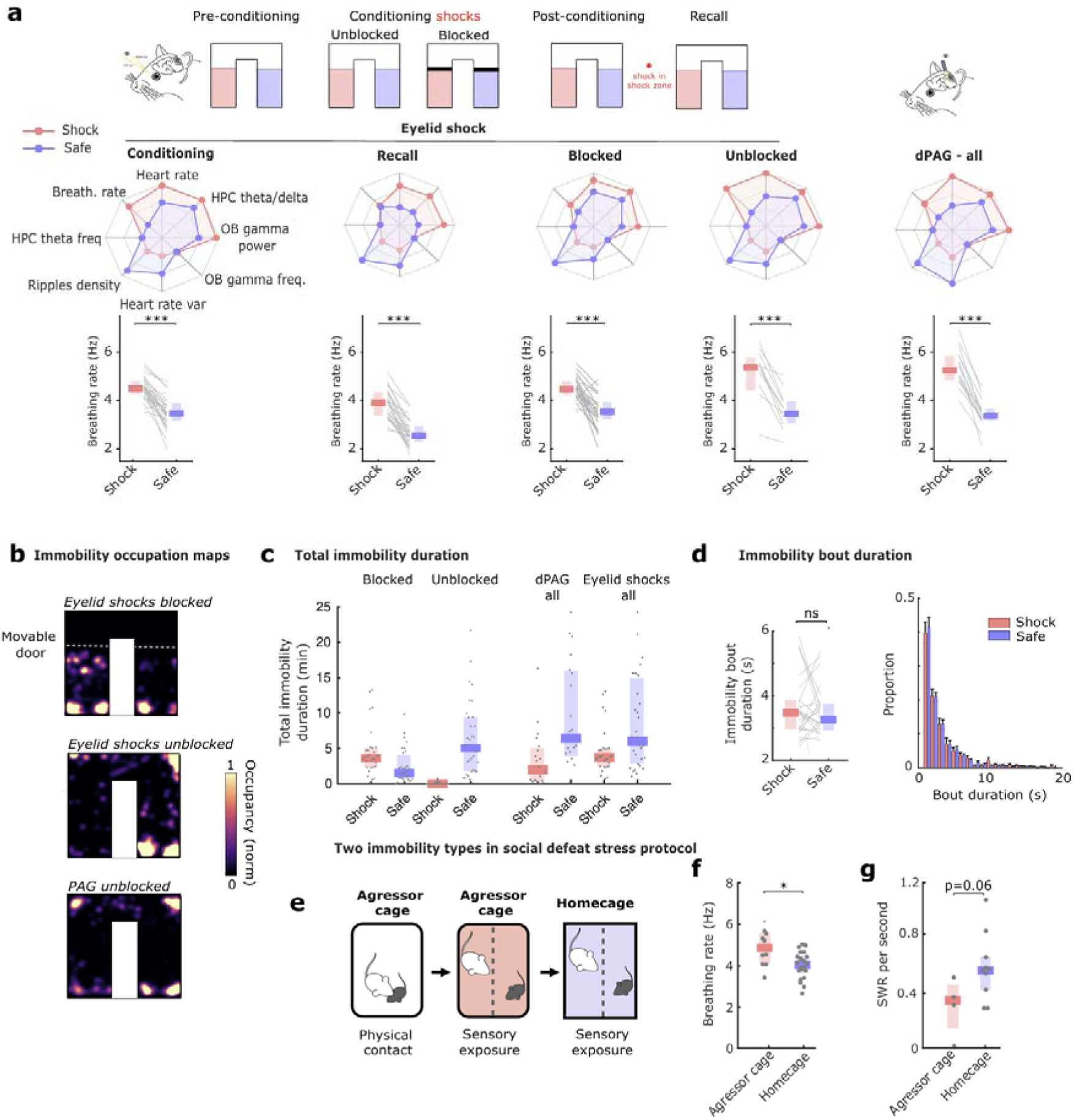
Robustness of two immobility states across conditions. a. Overview of the physiological characteristics of the shock and safe side immobility at different stages of the protocol and using dPAG aversive stimulation. For each condition, a spider plot gives the average value for each parameter showing that the characteristic profile is stable across conditions. The bottom row shows the breathing rate during both immobility types, showing that a robust statistical difference is always present (n=29 (eyelid), 20 (PAG) ***P=4.71 **×** 10^−6^, 1.22 **×** 10^−5^, 1.51 **×** 10^−5^, 9.76 **×** 10^−4^, 6.10 **×** 10^−5^, Wilcoxon signed rank test). b. Occupation maps showing where animals displayed immobility. Using eyelid shocks as an aversive stimulation, shock side immobility mainly occurred when animals were blocked by a door since the shock evoked a flight response. However shock side immobility was also spontaneously observed without the door (quantified in c). With dPAG stimulation, more shock side immobility was observed without the door because stimulation did not immediately induce flight. c. Total immobility duration in each condition. This shows that although rare, immobility for up to one minute does occur on the shock side in certain animals without being blocked using eyelid shocks. d. Immobility bout duration in shock and safe arms showing no difference between the two (n=29 (NS)P=0.6733, Wilcoxon signed rank test). e. Schematic of the social defeat protocol. The C57 mouse is first in contact with a large, aggressive CD1 male in the CD1 cage during which fighting occurs. After the encounter, the behaviour and physiology of the C57 mouse is recorded first in the CD1 cage and then in its own homecage. This allows us to contrast two environments with higher and lower levels of threat analogous to the shock and safe arms of the U-Maze. f. Breathing rate during immobility in the aggressor and home cage showing higher rates in the aggressor cage (n=10/21, *P=0.016, Wilcoxon rank sum test). g. SWR occurrence per second during immobility in the aggressor and home cage showing a tendency to a lower number of SWR in the aggressor cage. (n=8/4, *P=0.06, Wilcoxon rank sum test).

**Figure S3.**
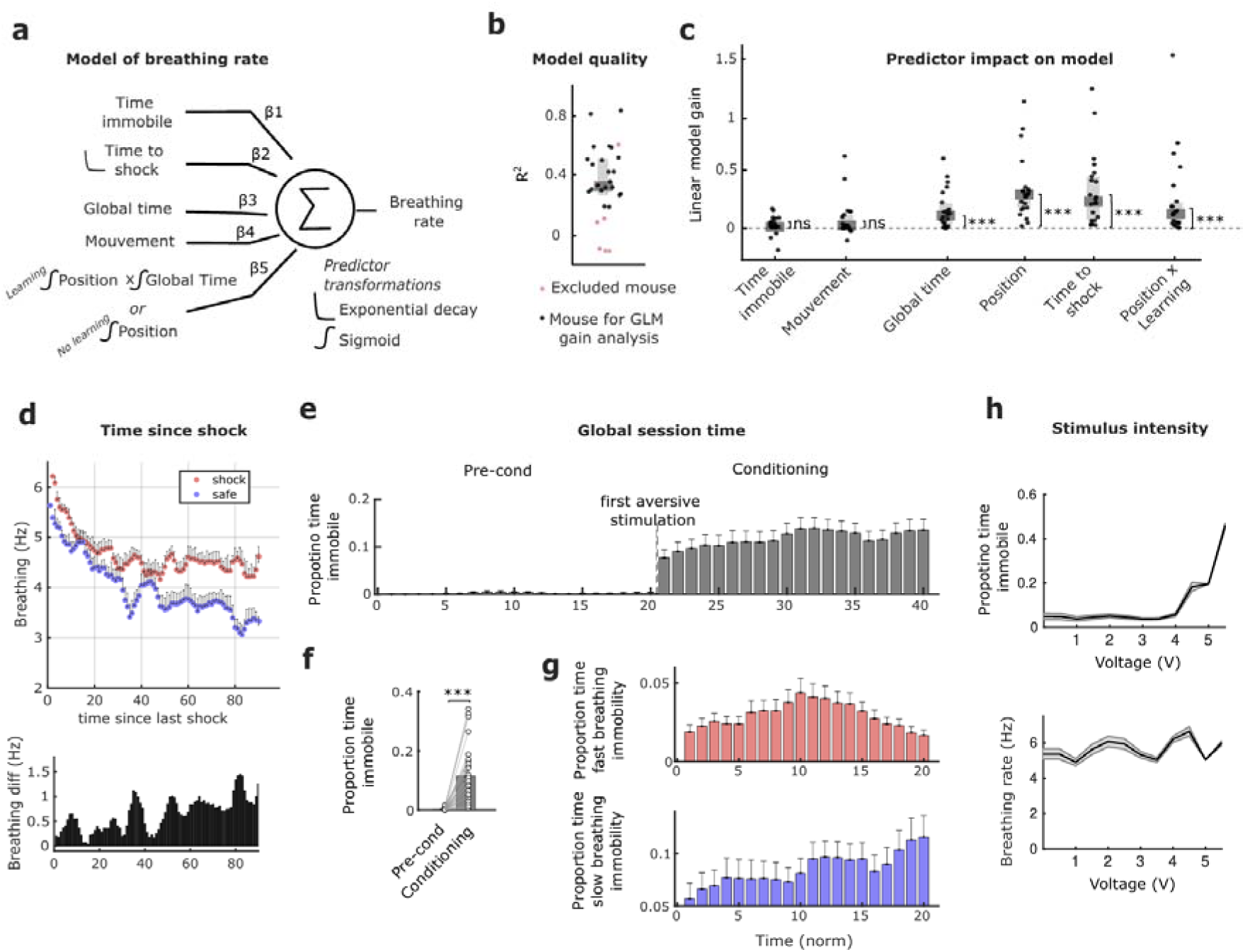
Robustness of two immobility states across conditions. a. Schematic showing the construction of the linear model that applies weights to each of the predictors in order to reconstruct breathing rate during immobility.^11^ b. R2 for the linear model fitted to each mouse after 5-fold cross validation. Mice shown in red were excluded from further analysis because they either had too little data or too poor quality of fit. c. Linear model gain for each predictor showing that neither the total time immobile nor the movement prior to immobility helped predict breathing rate. Global time in the protocol played a small role. The two strongest predictors are the time since the last shock and the position in the maze. Finally, the “learning” term included in the position predictor also plays a role in predicting breathing rate, showing that the difference between shock and safe side immobility emerges progressively. (n=22, NS= 0.11, 0.15, P=***P=3.3×10^−5^, 4.7×10^−7^, 4.7×10^−7^, 4.7×10^−7^, Wilcoxon rank sum test vs 0). d. Breathing rate as a function of the time since the mouse received the last shock during immobility on the safe or shock side. This shows a clear impact of shock time on breathing frequency, with an exponential decay. However the position of immobility has an impact at all time delays, with lower frequency observed in the safe arm. e. Proportion of time spent immobile throughout habituation and conditioning sessions, showing a sharp increase when mice receive the first shock. f. Mean proportion of time spent immobile during the habituation and conditioning (n=29, ***P=2.56 **×** 10^−5^, Wilcoxon signed rank test). g. Proportion of time spent immobile in the fast (top) or slow (bottom) breathing state. This shows that both states are found throughout the session although there is an increase in slow breathing immobility that we argue (Fig. 5) is linked to learning. h. Mice were submitted to unescapable eyelid shocks of increasing voltage that resulted in a progressive increase in time spent immobile without any corresponding change in breathing frequency. Thus, all stimulation strengths evoked freezing and not the putative recovery state.

**Figure S4.**
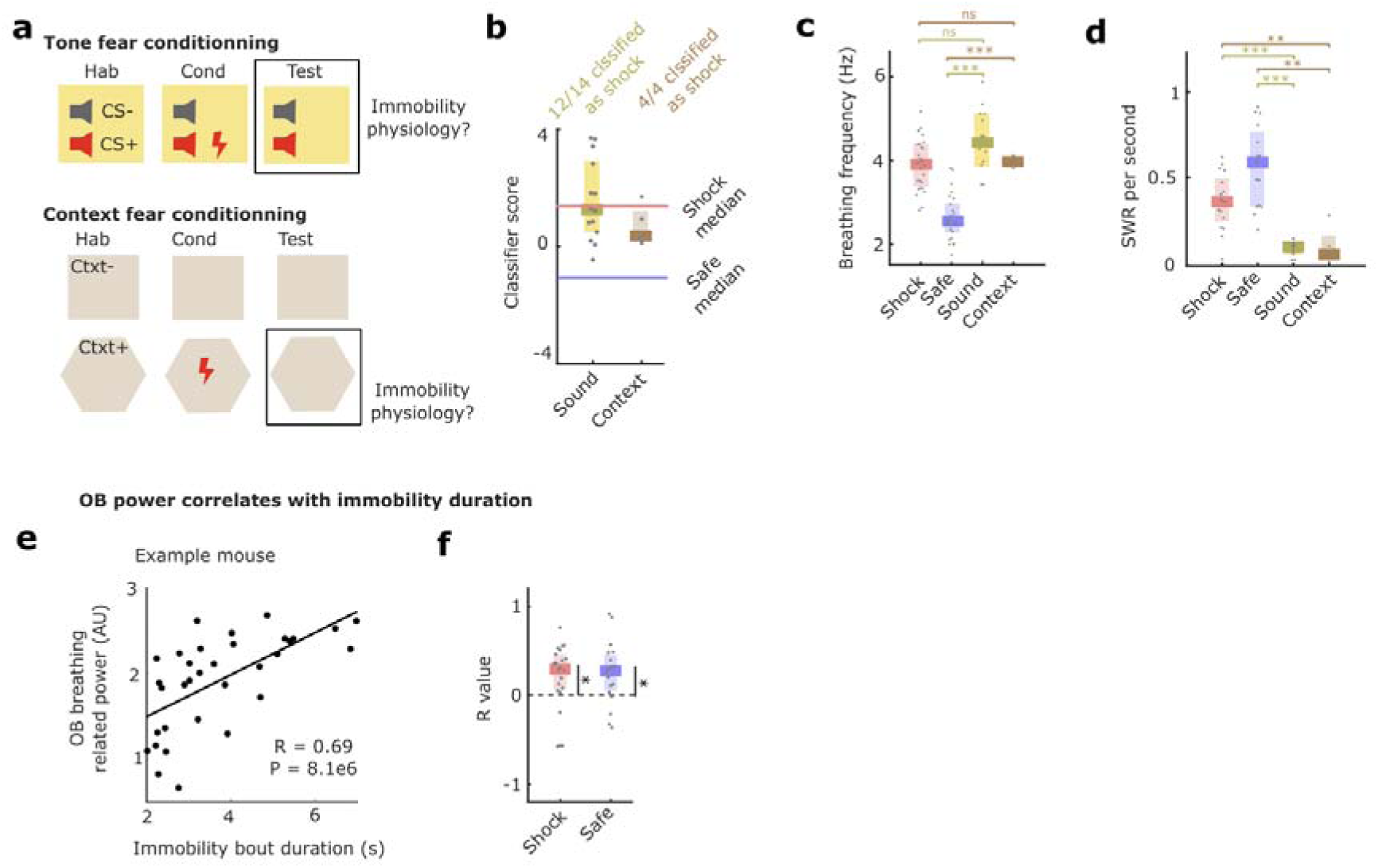
Shock side fast breathing immobility corresponds to freezing. a. Schematic showing the tone fear and contextual fear conditioning protocols. b. Classifier score for immobility (freezing) time during the two classical protocols showing that both are classified as being shock-like. c. Mean breathing frequency during shock and safe arm immobility, tone freezing and contextual freezing showing that breathing is similar during shock arm immobility and classical freezing (n=29/29/14/5, ***P=1.06 **×** 10^−4^, 9.70 **×** 10^−4^, (NS)P=0.4422, 0.5277, Wilcoxon rank sum test). d. SWR per second during shock and safe arm immobility, tone freezing and contextual freezing showing that SWR occurrence is low during shock arm immobility and classical freezing (n=29/29/14/5, comp. safe ***P=2.78 **×** 10^−5^, **P=0.0015, comp. shock ***P=1.23 **×** 10^−4^, **P=0.0077, Wilcoxon rank sum test). We had previously shown that 4 Hz breathing causally contributes to increasing freezing bouts duration and that stronger breathing power was linked to longer bouts. We verified this relation for the two immobility types we studied^11^. e. Data from an example mouse showing that longer immobility bouts are characterized by stronger breathing oscillations in the OB (n=33 freezing episodes of a mouse, R=0.6922, P=8.0848 **×** 10^−6^, Spearman correlation). f. Correlation coefficient for all mice during shock and side immobility between immobility bout duration and oscillation power. This shows that in both states there is a positive correlation (n=29, comp. safe *P=0.049, 0.011, Wilcoxon rank sum test).

**Figure S5.**
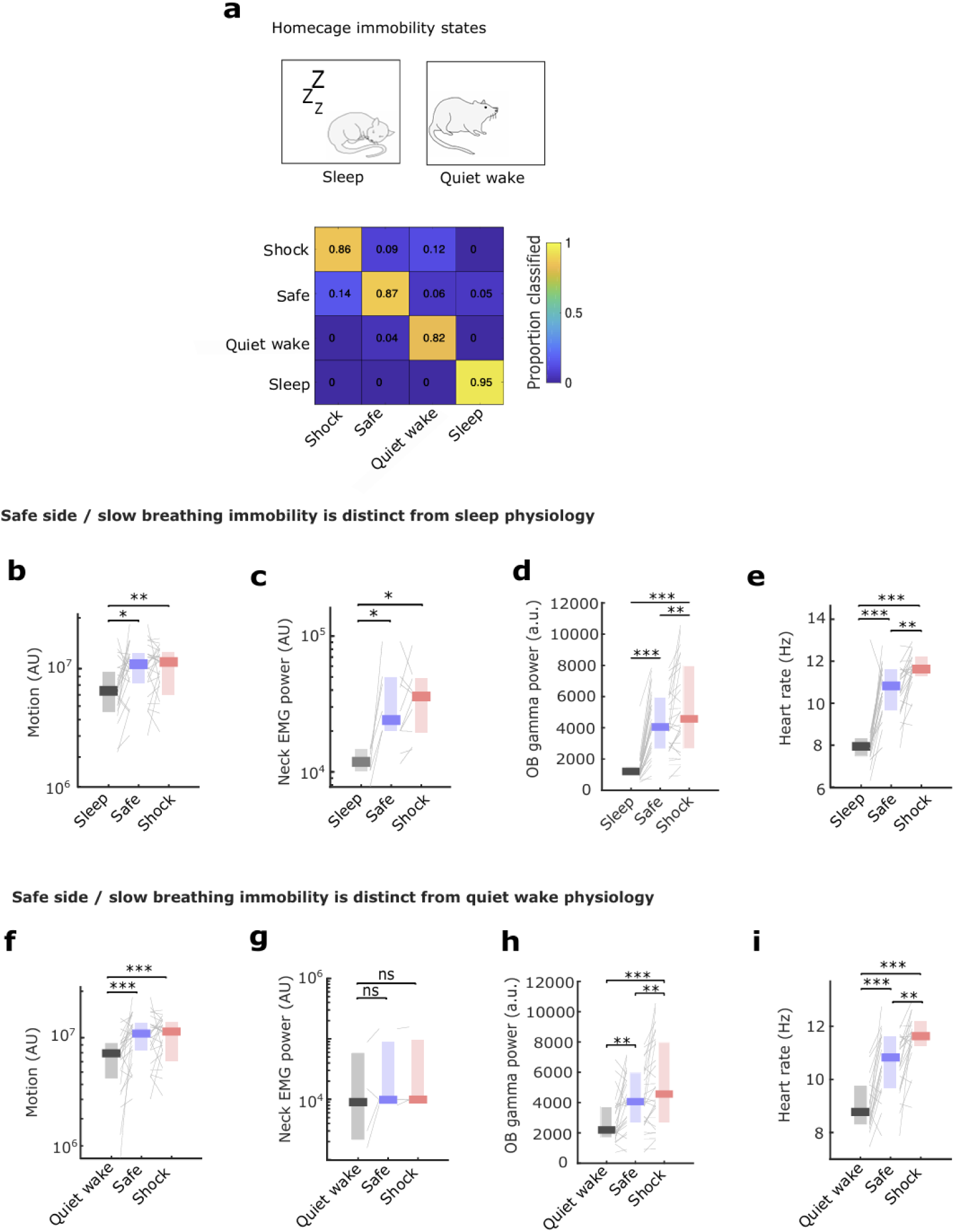
Safe side slow breathing immobility is distinct from sleep and quiet wake. a. During preconditioning homecage sessions we characterized the physiology of quiet wakefulness and sleep. We then trained a classifier to discriminate between these states and the shock and safe-arm immobility. The classifier was trained using a leave-one-out cross-validation approach: for each mouse (n), the model was trained on the data from the remaining (n–1) mice, and the resulting classifier was used to predict the immobility score for the held-out mouse on both the shock and safe sides. This procedure was repeated for every mouse. The confusion matrix shows that all four states can be clearly distinguished, without confusion between safe arm immobility and non-aversive related quiescent states. b-e. Mean motion (measured by head acceleration), nuchal EMG, heart rate, OB gamma power and heart rate for sleep, shock and safe side immobility showing that all variables are significantly different (b. **P=0.0072, *P=0.0177, c. *P=0.0312, 0.0312, d. ***P=3.7896 **×** 10^−6^, 2.5631 **×** 10^−6^, e. ***P=1.2042 **×** 10^−4^, 1.6286 **×** 10^−4^, Wilcoxon signed rank test). f-i. Mean motion (measured by head acceleration), nuchal EMG, heart rate, OB gamma power and heart rate for quiet wake, shock and safe side immobility showing that all variables except EMG are significantly different (f. ***P=5.2571 **×** 10^−4^, 5.4594 **×** 10^−4^, g. (NS)P=0.0569, 0.2969, h. ***P=1.7735 **×** 10^−5^, 2.4254 **×** 10^−4^, i. 1.5510 **×** 10^−4^, 3.9814 **×** 10^−4^, Wilcoxon signed rank test)..

**Figure S6.**
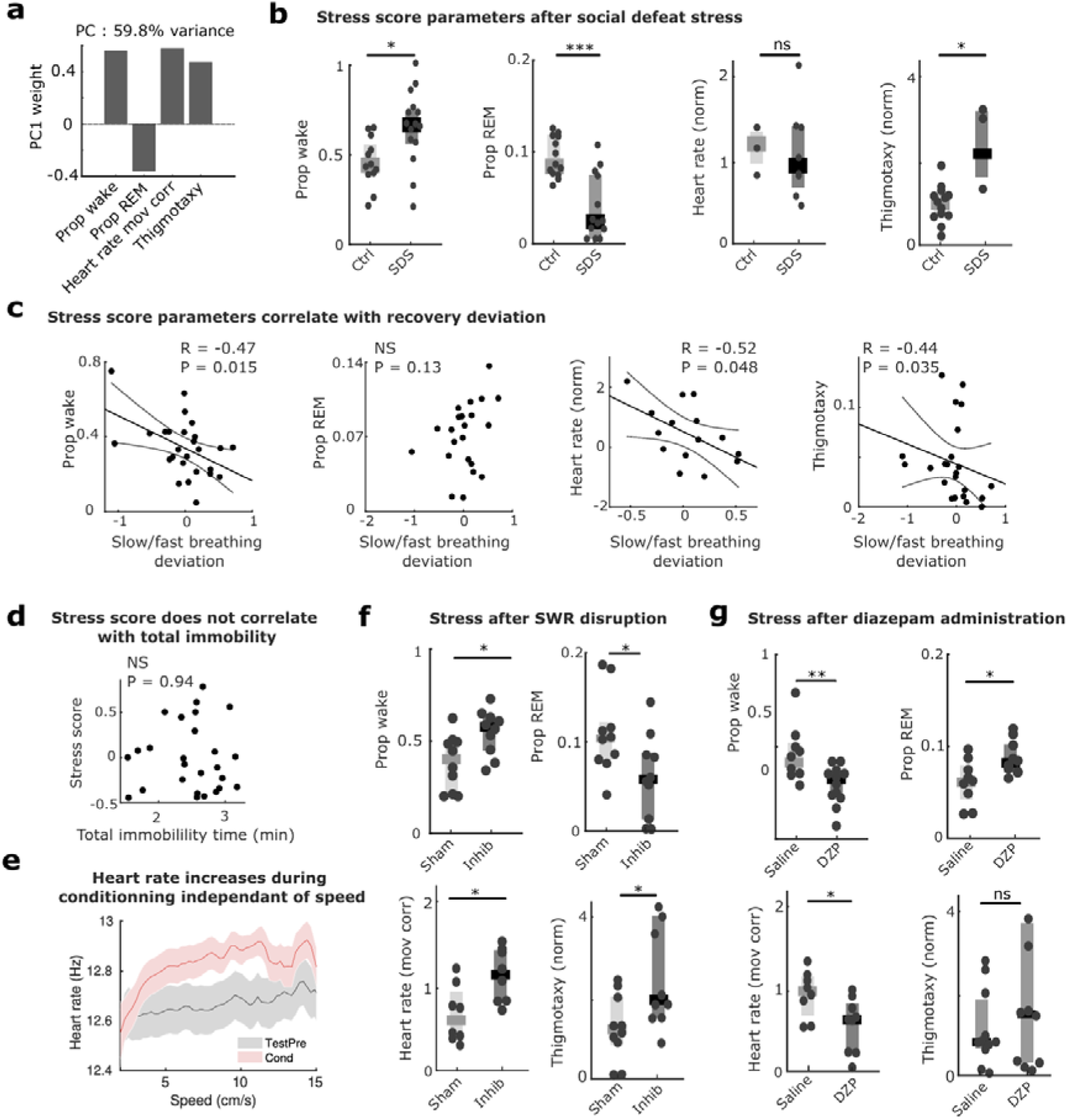
Details of stress score individual parameters. a. Weights on the first principal component that explained 60% of variance across mice for the four individual parameters compounded by the stress score. Note that each component has a similar contribution. b. The components of the stress score, except for heart rate during movement, all increased after animals were submitted to social defeat stress (n=13/14, 13/14, 13/8, 4/3, *P=0.01, 0.0007, 0.93, 0.014, Wilcoxon rank sum test). c. The components of the stress score, except for REM proportion, all correlated with the recovery deviation measured during U-Maze conditioning. d. The stress score did not correlate with total time spent immobile in the U-Maze conditioning. e. Heart rate as a function of speed during preconditioning and conditioning sessions showing that conditioning increases heart rate for all speeds. Error bars are SEM (n=22). f. The components of the stress score all indicated an increase in stress after SWR disruption (n=10/10, *P=0.02, 0.04, 0.02, 0.03 Wilcoxon rank sum test). g. The components of the stress score, except for thigmotaxis, all indicated a decrease in stress after SWR disruption (n=10/10, *P=0.0095, 0.028, 0.04, 0.35, Wilcoxon rank sum test).

**Figure S7.**
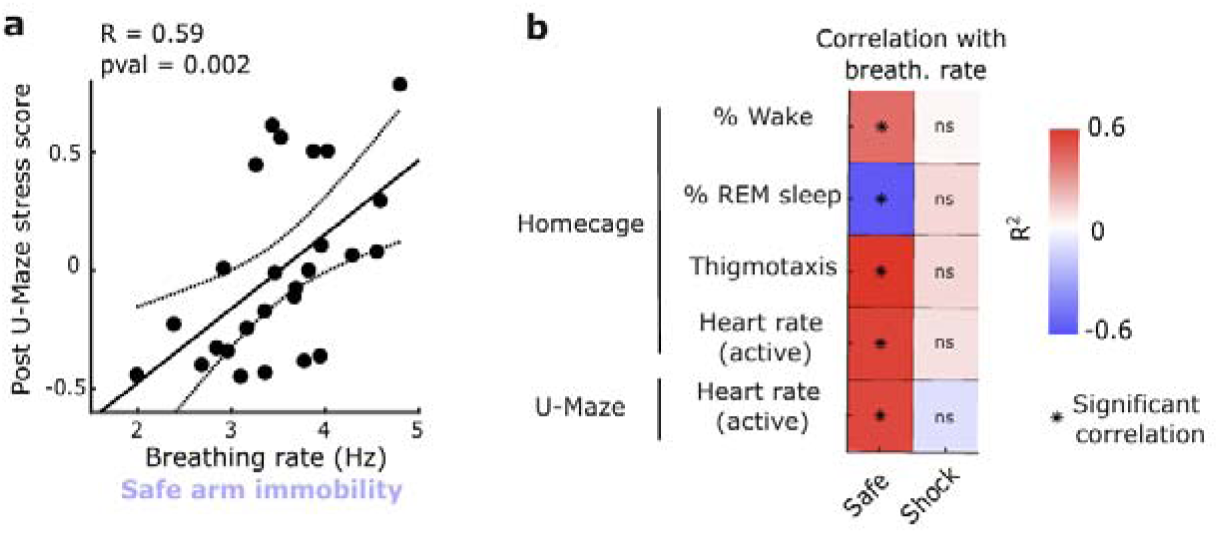
Safe side immobility characteristics predict post task recovery. a. Correlation between breathing rate during safe side immobility and the post-task stress score (n=29, R=0.5904, P=0.0018, Spearman correlation). b. Correlation coefficient indicated by color between breathing rate during safe or shock side immobility and all elements of the post-task stress score as well as heart rate during the task that we use as an indicator for stress during the task. This shows that safe side but not shock side characteristics are predictive of stress during and after the task (safe: R=0.4448, -0.5675, 0.5474, 0.6000, 0.5157, P=0.0238, 0.0067, 0.0077, 0.0199, shock: R=-0.0869, 0.0593, 0.0999, 0.1929, -0.0271, P=0.6786, 0.7934, 0.6572, 0.4901, 0.9114).

**Figure S8.**
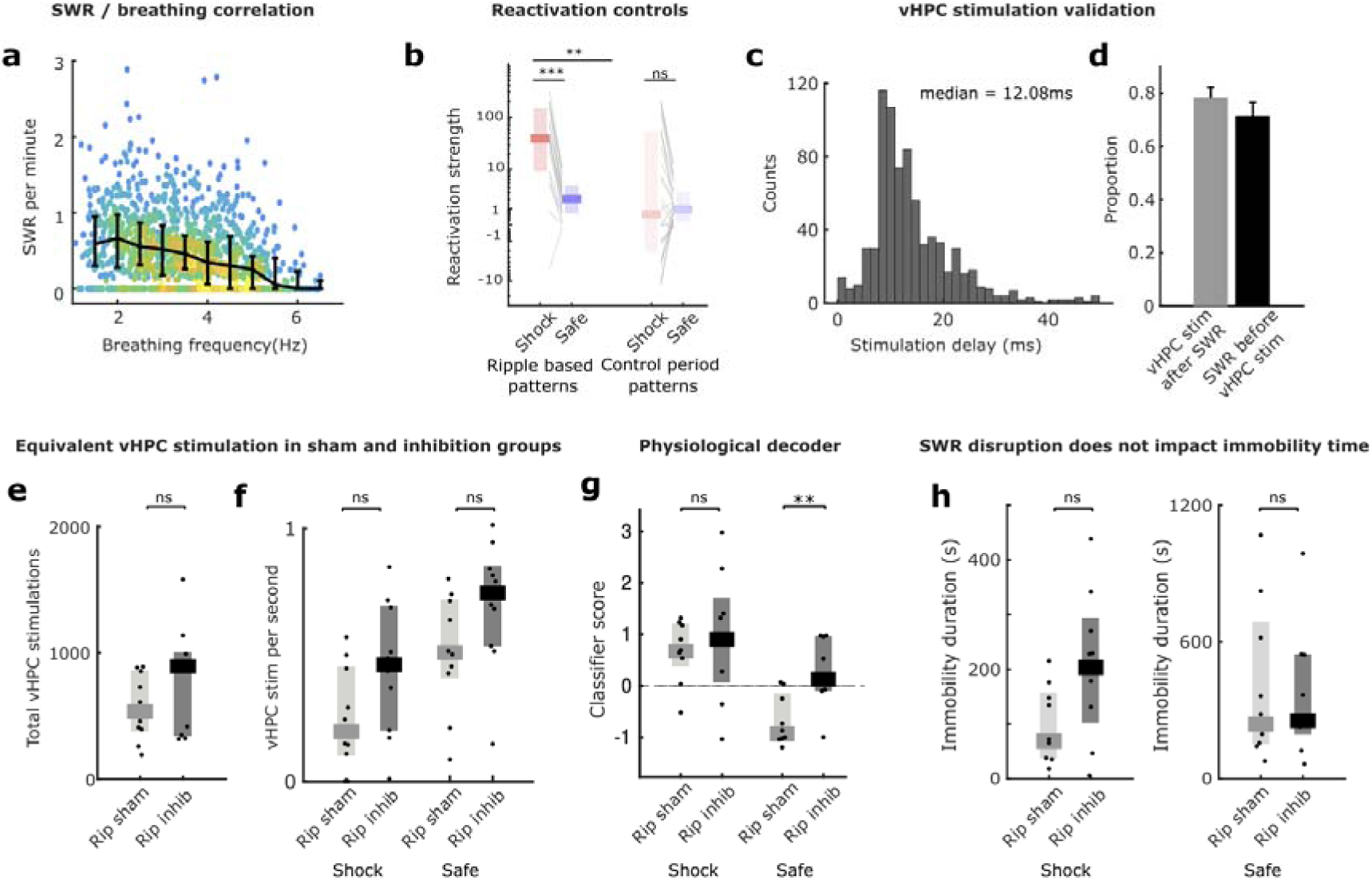
Controls for SWR disruption. a. Correlation between instantaneous (2s bins) SWR occurrence and breathing rate across all animals during all immobility periods. This shows that the lower the breathing, the more likely SWRs are to occur (R=-0.30, P=-2.48 **×** 10^−27^, Pearson correlation). b. Mean activation strength of all ripple-identified patterns during shock or safe side exploration (left) and of all pre-ripple identified patterns, used as a control period (right). Note that activation strength is lower and shows no difference between shock and safe side, indicating that preferential reactivation of shock side activity is specific to SWRs periods. c. Distribution of delay between an online detected SWRs and the stimulation of the vHPC in ripple disrupted animals. d. Specificity and selectivity of the vHPC stimulation relative to offline detected SWRs was around 80%. e-f. Total number and density during immobility of vHPC stimulations in control and disruption groups showing no difference (n=10/10, (NS)P=0.2123, 0.0889, Wilcoxon rank sum test). g. Classifier score of immobility in shock and safe arms showing that shock arm immobility was well classified in both groups but safe arm immobility in ripple disruption group tended to be classified as more shock-like (n=10/10, (NS)P=0.6250, **P=0.002, Wilcoxon rank sum test). h. Immobility duration in shock (left) and safe (right) arms showing no difference between control and disruption groups (n=10/10, (NS)P=0.0757, 0.9698, Wilcoxon rank sum test)..

**Figure S9.**
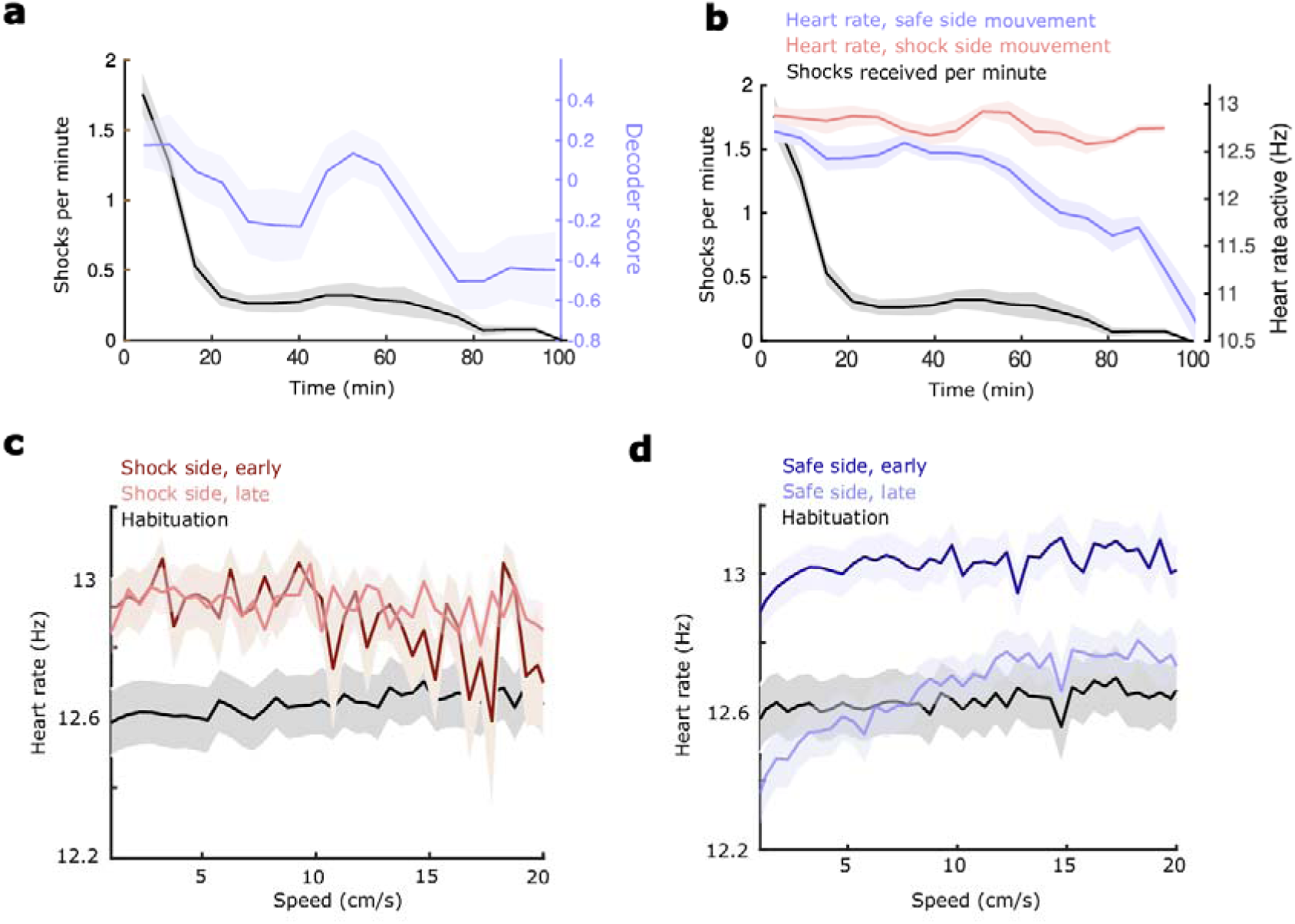
Safety learning indicated by safe side immobility and heart rate. a. Temporal evolution through conditioning of safe side immobility classifier score (blue), and shock number (gray). This shows that avoidance learning, indexed by shock density, occurs faster than safety learning, indexed by the shift in global physiology during safe side immobility to the slow breathing type (n=29, error bars are s.e.m.). b. Temporal evolution through conditioning of heart rate during movement on the safe side (blue), the shock side (red) and shock number (gray). This shows that heart rate remains high on the shock side throughout conditioning but gradually declines during the task on the safe side. This decline in heart rate on the safe side can be taken as indicative of the mouse learning that this is a non threatening region. c. Average heart rate at different speeds during habituation and during the beginning or end of conditioning in the shock zone. This shows that throughout conditioning, heart rate is higher in the shock zone than during habituation, irrespective of speed, consistent with it indicating the stress related to the aversive zone of the task. d. Average heart rate at different speeds during habituation and during the beginning or end of conditioning in the safe zone. This shows that early in the task heart rate is increased in the safe zone relative to a non-stressful condition irrespective of speed but then progressively drops, consistent with the animal learning that the safe zone is a less stressful environment.

**Figure S10.**
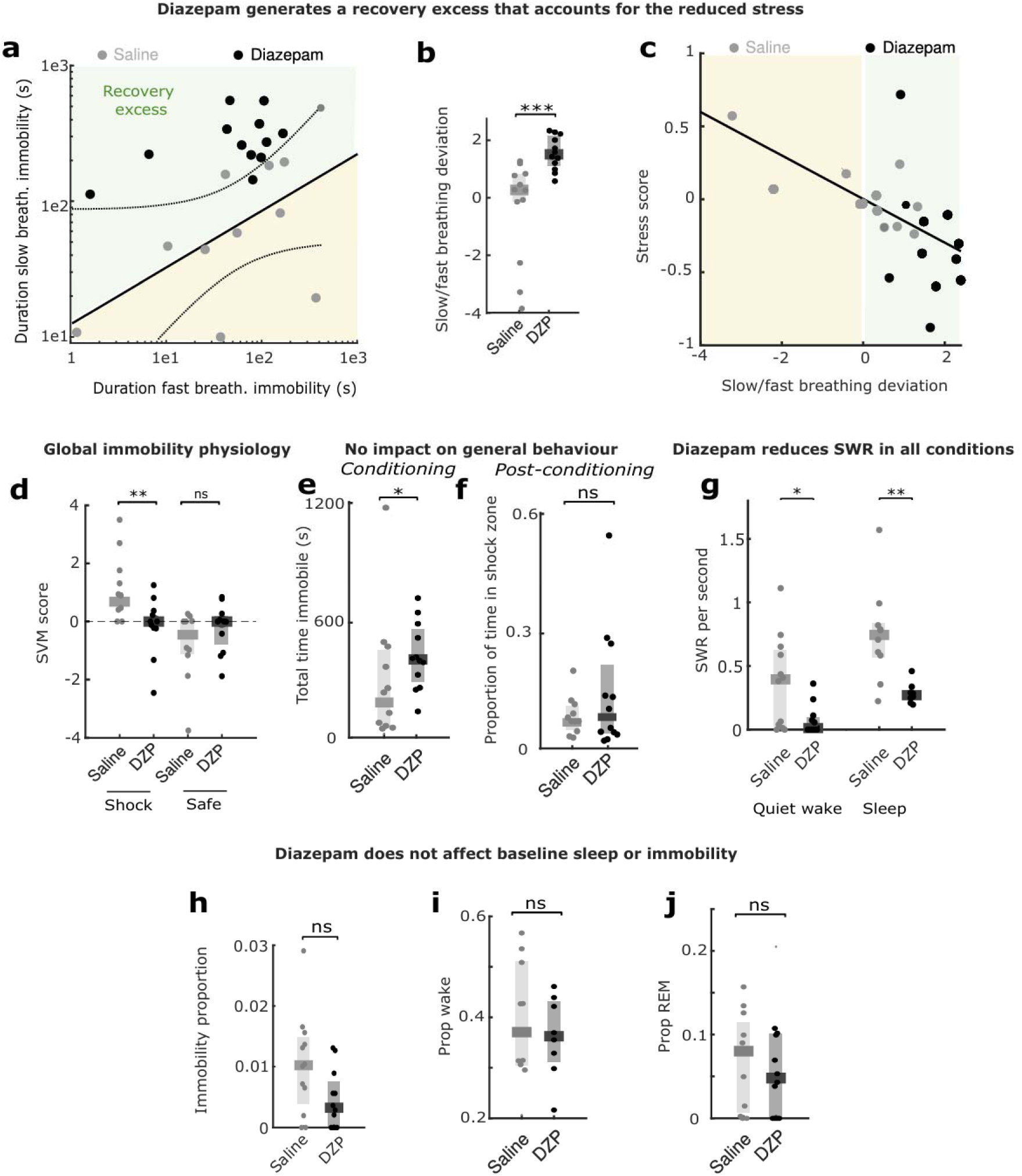
Controls for diazepam administration. a. Time in the slow breathing immobility state is strongly correlated with time in the slow breathing immobility state in the saline group. The diazepam group is clearly shifted towards an increase in slow breathing states, which we quantify with the slow breathing deviation relative to the relationship defined using saline mice. b. Slow/fast breathing deviation for saline and diazepam groups.(n=11/12, *P=4 **×** 10^−4^, Wilcoxon rank sum test). c. The slow/fast breathing deviation is correlated with the post-task stress score across the two groups. d. Classifier score of immobility in shock and safe arms showing that safe arm immobility was well classified in both groups but shock arm immobility in the diazepam group tended to be classified as more safe-like (n=11/12, **P=0.0055, (NS)P=0.3213, Wilcoxon rank sum test). e. Total time spent immobile was not affected by diazepam (n=11/12, *P=0.045, Wilcoxon rank sum test). f. The proportion of time spent in the shock zone post-conditioning was not affected by diazepam. g. Diazepam reduces SWR occurrence during sleep and quiet wake (n=11/12, (NS)P=0.8294, Wilcoxon rank sum test). h-j. When diazepam is administered prior to a homecage session it did not impact proportion of time spent immobile (h), the breathing during these immobile periods (i) or sleep architecture (proportion wake (j) or in REM (k)) that we used as stress markers previously (n=11/12, h. (NS)P=0.0513, j. (NS)P=0.8286, k. (NS)P=0.5413, Wilcoxon rank sum test).

## References

1. Fanselow, Michael S & Lester, Laurie S. A functional behaviouristic approach to aversively motivated behaviour: predatory imminence as a determinant of the topography of defensive behaviour. in Evolution and learning. (1988).

2. Mobbs, D. The ethological deconstruction of fear(s). Curr. Opin. Behav. Sci. 24, 32–37 (2018).

3. Apfelbach, R., Blanchard, C. D., Blanchard, R. J., Hayes, R. A. & McGregor, I. S. The effects of predator odors in mammalian prey species: A review of field and laboratory studies. Neurosci. Biobehav. Rev. 29, 1123–1144 (2005).

4. Sukikara, M. H., Mota-Ortiz, S. R., Baldo, M. V., Felicio, L. F. & Canteras, N. S. The periaqueductal gray and its potential role in maternal behaviour inhibition in response to predatory threats. Behav. Brain Res. 209, 226–233 (2010).

5. Bolles, R. C. & Fanselow, M. S. A perceptual-defensive-recuperative model of fear and pain. Behav. Brain Sci. 3, 291–301 (1980).

6. Sapolsky, R. M. Stress and the brain: individual variability and the inverted-U. Nat. Neurosci. 18, 1344–1346 (2015).

7. Dong, Y. et al. Stress relief as a natural resilience mechanism against depression-like behaviours. Neuron 111, 3789–3801.e6 (2023).

8. Yu, X. et al. A specific circuit in the midbrain detects stress and induces restorative sleep. Science 377, 63–72 (2022).

9. Troisi, A. Displacement Activities as a behavioural Measure of Stress in Nonhuman Primates and Human Subjects. Stress 5, 47–54 (2002).

10. Kalueff, A. V. et al. Neurobiology of rodent self-grooming and its value for translational neuroscience. Nat. Rev. Neurosci. 17, 45–59 (2016).

11. Bagur, S. et al. Breathing-driven prefrontal oscillations regulate maintenance of conditioned-fear evoked freezing independently of initiation. Nat. Commun. 12, 2605 (2021).

12. Jhang, J., Park, S., Liu, S., O’Keefe, D. D. & Han, S. A top-down slow breathing circuit that alleviates negative affect in mice. Nat. Neurosci. 27, 2455–2465 (2024).

13. De Sousa Abreu, R. P., et al. Episodic slow breathing in mice markedly reduces fear responses. Preprint at 10.1101/2024.12.09.627565 (2024).

14. Ma, D. et al. Benefits From Different Modes of Slow and Deep Breathing on Vagal Modulation. IEEE J. Transl. Eng. Health Med. 12, 520–532 (2024).

15. Papez, J. W. A PROPOSED MECHANISM OF EMOTION. Arch. Neurol. Psychiatry 38, 725 (1937).

16. Gray, J. A. The neuropsychology of anxiety: An enquiry into the functions of the septo-hippocampal system. Behav. Brain Sci. 5, 469–484 (1982).

17. O’Keefe, J. & Nadel, L. The hippocampus as a cognitive map. Behav. Brain Sci. 2, 487–494 (1979).

18. Buckner, R. L. The Role of the Hippocampus in Prediction and Imagination. Annu. Rev. Psychol. 61, 27–48 (2010).

19. Wu, C.-T., Haggerty, D., Kemere, C. & Ji, D. Hippocampal awake replay in fear memory retrieval. Nat. Neurosci. 20, 571–580 (2017).

20. Calvin, O. L., Erickson, M. T., Walters, C. J. & Redish, A. D. Dorsal hippocampus represents locations to avoid as well as locations to approach during approach-avoidance conflict. PLOS Biol. 23, e3002954 (2025).

21. Appelhans, B. M. & Luecken, L. J. Heart Rate Variability as an Index of Regulated Emotional Responding. Rev. Gen. Psychol. 10, 229–240 (2006).

22. Vianna, D. M. L. & Carrive, P. Changes in cutaneous and body temperature during and after conditioned fear to context in the rat. Eur. J. Neurosci. 21, 2505–2512 (2005).

23. Bagur, S. et al. Harnessing olfactory bulb oscillations to perform fully brain-based sleep-scoring and real-time monitoring of anaesthesia depth. PLoS Biol. 16, e2005458 (2018).

24. Moberly, A. H. et al. Olfactory inputs modulate respiration-related rhythmic activity in the prefrontal cortex and freezing behaviour. Nat. Commun. 9, 1528 (2018).

25. Carrive, P. The periaqueductal gray and defensive behaviour: Functional representation and neuronal organization. Behav. Brain Res. 58, 27–47 (1993).

26. Pawlyk, A. C., Morrison, A. R., Ross, R. J. & Brennan, F. X. Stress-induced changes in sleep in rodents: Models and mechanisms. Neurosci. Biobehav. Rev. 32, 99–117 (2008).

27. Rosso, M. et al. Reliability of common mouse behavioural tests of anxiety: A systematic review and meta-analysis on the effects of anxiolytics. Neurosci. Biobehav. Rev. 143, 104928 (2022).

28. Stiedl, O., Jansen, R. F., Pieneman, A. W., Ögren, S. O. & Meyer, M. Assessing aversive emotional states through the heart in mice: Implications for cardiovascular dysregulation in affective disorders. Neurosci. Biobehav. Rev. 33, 181–190 (2009).

29. Buzsáki, G. Hippocampal sharp wavelripple: A cognitive biomarker for episodic memory and planning. Hippocampus 25, 1073–1188 (2015).

30. Girardeau, G., Benchenane, K., Wiener, S. I., Buzsáki, G. & Zugaro, M. B. Selective suppression of hippocampal ripples impairs spatial memory. Nat. Neurosci. 12, 1222–1223 (2009).

31. Oliva, A., Fernández-Ruiz, A., Leroy, F. & Siegelbaum, S. A. Hippocampal CA2 sharp-wave ripples reactivate and promote social memory. Nature 587, 264–269 (2020).

32. Roux, L., Hu, B., Eichler, R., Stark, E. & Buzsáki, G. Sharp wave ripples during learning stabilize the hippocampal spatial map. Nat. Neurosci. 20, 845–853 (2017).

33. Ma, X. et al. The Effect of Diaphragmatic Breathing on Attention, Negative Affect and Stress in Healthy Adults. Front. Psychol. 8, 874 (2017).

34. Lehrer, P. M. & Gevirtz, R. Heart rate variability biofeedback: how and why does it work? Front. Psychol. 5, (2014).

35. Yackle, K. et al. Breathing control center neurons that promote arousal in mice. Science 355, 1411–1415 (2017).

36. Solomon, R. L. The opponent-process theory of acquired motivation: The costs of pleasure and the benefits of pain. Am. Psychol. 35, 691–712 (1980).

37. Girardeau, G., Inema, I. & Buzsáki, G. Reactivations of emotional memory in the hippocampus– amygdala system during sleep. Nat. Neurosci. 20, 1634–1642 (2017).

38. Pedraza, L. K. et al. Hippocampal sharp wave ripples mediate generalization and subsequent fear attenuation via closed-loop brain stimulation in rats. Preprint at 10.1101/2024.04.30.591894 (2024).

39. Schiller, D. et al. Preventing the return of fear in humans using reconsolidation update mechanisms. Nature 463, 49–53 (2010).

40. Sevinc, G. & Lazar, S. W. How does mindfulness training improve moral cognition: a theoretical and experimental framework for the study of embodied ethics. Curr. Opin. Psychol. 28, 268–272 (2019).

41. Roelofs, K. Freeze for action: neurobiological mechanisms in animal and human freezing. Philos. Trans. R. Soc. B Biol. Sci. 372, 20160206 (2017).

42. Tomizawa, H. et al. A Transient Fear Reduction by Pair-Exposure with a Non-Fearful Partner during Fear Extinction Independent from Corticosterone Level in Mice. J. Behav. Brain Sci. 03, 415–421 (2013).

43. Dopfel, D. et al. Individual variability in behaviour and functional networks predicts vulnerability using an animal model of PTSD. Nat. Commun. 10, 2372 (2019).

44. Malikowska-Racia, N., Salat, K., Gdula-Argasinska, J. & Popik, P. Sex, Pramipexole and Tiagabine Affect behavioural and Hormonal Response to Traumatic Stress in a Mouse Model of PTSD. Front. Pharmacol. 12, 691598 (2021).

45. Rogan, M. T., Leon, K. S., Perez, D. L. & Kandel, E. R. Distinct Neural Signatures for Safety and Danger in the Amygdala and Striatum of the Mouse. Neuron 46, 309–320 (2005).

46. Rojas-LÃ-bano, D., Frederick, D. E., EgaÃ±a, J. I. & Kay, L. M. The olfactory bulb theta rhythm follows all frequencies of diaphragmatic respiration in the freely behaving rat. Front. Behav. Neurosci. 8, (2014).

47. Peyrache, A., Khamassi, M., Benchenane, K., Wiener, S. I. & Battaglia, F. P. Replay of rule-learning related neural patterns in the prefrontal cortex during sleep. Nat. Neurosci. 12, 919–926 (2009).

