## Supplementary material for "Hippocampal reactivation of aversive experience enables safety learning and slow-breathing state for recovery from stress": Stats supplementary

| Fig1.b, pre conditioning | Wilcocxon signed rank | n=29 | p=0.3695 zval=0.8974 |
| --- | --- | --- | --- |
| Fig1.b, post conditioning | Wilcocxon signed rank | n=29 | p=5.321 **×** 10^-6^  zval=0.8974 |
| Fig1.e | Wilcocxon signed rank | n=28 | p=9.7817 **×** 10^-6^ zval=-4.4219 |
| Fig1.f | Wilcocxon signed rank | n=23 | Moving-shock  p=2.7016 **×** 10^-5^ zval=4.1973  Moving-safe  p=1.8215 **×** 10^-5^ zval=4.2857  shock-safe  p=0.9273  zval=0.0912 |
| Fig1.j | Wilcocxon signed rank | n= 28 | p=4.7157 **×** 10^-6^ zval=4.5771 |
| Fig1.k | Wilcocxon signed rank | n= 22 | p=2.0135 **×** 10^-4^ zval=3.7173 |
| Fig1.l | Wilcocxon signed rank | n= 22 | p=3.3392 **×** 10^-4^ zval=-3.5875 |
| Fig. 2. b | Wilcoxon rank-sum | n=13, n=14 sessions | p=0.0004 Zval=-3.5 |
| Fig. 2. c | Pearson correlation | n=20 | R=0.71157  p=9.2719 **×** 10^-4^ |
| Fig. 2.d | Pearson correlation  Wilcocxon signed rank | n=28 | R=0.66292  p=8.9032 **×** 10^-5^  Boxplot  P=2.5631 **×** 10^-6^  zval=-4.7030 |
| Fig. 2e | Spearman correlation | n=26 | R=-0.50632  p=0.0090898 |
| Fig. 3b | Wilcocxon signed rank | n= 19 | p=1.3183 **×** 10^-4^ zval=-3.8230 |
| Fig. 3e | Wilcoxon rank sum | n=21, n=82, n=54 | shock/middle  p=0.0118  zval=2.5171  shock/safe  p=5.3367 **×** 10^-4^  zval=3.4633 |
| Fig. 3i | Wilcocxon signed rank | n=22 , n= 22 | p=5.300 **×** 10^-5^ zval=4.042 |
| Fig. 4b | Wilcoxon rank sum | shock  n=8 , n= 8  safe  n=7, n=9 | shock  p=0.3961 stat=59.5  safe  p=0.0023 stat=32 |
| Fig. 4.c | Wilcoxon rank sum | n=10, n=10 | p=0.0046  zval=-2.8347 |
| Fig. 4.d, sham | Spearman correlation | n=10 | R=0.68; P=0.049 |
| Fig. 4.d, Rip inhib | Spearman correlation | n=10 | R=0.72; P=0.024 |
| Fig. 4. e | Wilcoxon rank sum | n=8/8 | p=0.0093  ranksum=78 |
| Fig. 4.f, Rip inhib + sham | Spearman correlation | n=16 | R=-0.39; P=0.089 |
| Fig. 4. g | Wilcoxon rank sum | n=10, n=10 | p=0.7337  zval=0.3402 |
| Fig. 4. i | Wilcoxon rank sum | n= 10 , n= 10 | p=0.7337 stat=0.3402 |
| Fig. 5. b | Wilcocxon signed rank | shock, n=28  safe, n=29 | shock  p=0.9637  zval=-0.0455  safe  p=0.0042  zval=2.8651 |
| Fig. 5. c | Wilcocxon signed rank | shock, n=28  n=29 | shock  p=0.6325  zval=-0.4782  safe  p=0.0017  zval=-3.1462 |
| Fig. 5. f |  | n=9, n=10 | rip sham - rip inhib  p=0.0312  zval=-2.1544  rip sham - 0  p=0.0098  signedrank=3  rip inhib - 0  p=0.3750  signedrank=18 |
| Fig. 6. b | Wilcoxon rank-sum | n=11, n=12 | fast  p=0.8294  zval=0.2154  slow  p=0.0074  zval=-2.6772 |
| Fig. 6. c | Wilcoxon rank-sum | n=11, n=12 | p=0.0392  zval=2.0618 |
| Fig. 6. e | Wilcoxon rank-sum | n=11, n=12 | Saline - Diazepam  p=0.0905  zval=-1.6925  Saline - 0  p=0.0098  signedrank=5  Diazepam - 0  p=0.3394  signedrank=26 |
| Fig. 6. f | Wilcoxon rank-sum | n=11, n=12 | shock  p=0.0121  ranksum=119  safe  p=0.5248  zval=0.6360 |
| Fig. 6. g | Wilcoxon rank-sum |  | shock  p=0.0019  ranksum=96  safe  p =5.7589 **×** 10^-4^  ranksum=105 |
| Supplementary figures | | | |
| Fig. S1a | Wilcoxon signed rank vs 0 | n=28, 22, 22, 22, 20, 27, 28, 27, 28, 27, 7 | p=7.2e-06, 2.59e-04, 1.76e-04, 8.75e-04, 8.91e-04, 2.91e-05, 6.91e-04, 0.91, 0.24, 0.17, 0.21  stats=4.48, 3.65, -3.74, 3.32, -3.3226, 4.18, 3.39, 0.10, 1.16, -1.37, 6 |
| Fig. S1b | Wilcocxon signed rank | n=8 | p=0.0391 signedrank=3 |
| Fig. S1d, gamma power | Wilcocxon signed rank | n=28 | p=0.0015  zval=3.1652 |
| Fig. S1e, gamma frequency | Wilcocxon signed rank | n=28 | p=0.0019  zval=3.1123 |
| Fig. S1g, theta/delta ratio | Wilcocxon signed rank | n=27 | p=2.618 **×** 10^-5^  zval=4.2044 |
| Fig. S1i, SWR power | Wilcocxon signed rank | n=26 | p=6.3294 **×** 10^-5^  zval=-4.002 |
| Fig. S1k, SVM | Wilcocxon signed rank | n=22 | p=4.6125 **×** 10^-5^  zval=4.0744 |
| Fig. S2a | Wilcocxon signed rank | Eyelid n= 28  PAG n=20  Unblocked=11 | Cond p=4.7157 **×** 10^-6^ zval=4.5771  Recall  p=1.2290 **×** 10^-5^  zval=4.3724  PAG  p=6.1035 **×** 10^-5^  signedrank=120  Blocked  p=1.5145 **×** 10^-5^  zval=4.3266  Unlocked  p=9.7656 **×** 10^-4^  signedrank=66 |
| Fig. S2c | Wilcoxon signed rank | Eyelid n= 28  PAG n=20 | Blocked  p=0.0042  zval=2.8611  Unblocked  p=1.1742 **×** 10^-6^  zval=-4.8599  PAG  p=0.0045  zval=-2.8373  Eyelid  p=0.0013  zval=-3.2138 |
| Fig. S2d | Wilcoxon signed rank | n= 28 | p=0.6733  zval=-0.4217 |
| Fig. S2f |  | n= 10, 21 control or SDS sessions | p=0.016, statistic=2.38 |
| Fig. S2g |  | n= 8, 4 control or SDS sessions | p=0.06, statistic=17 |
| Fig. S3f | Wilcoxon signed rank | n=28 | p=2.5631 **×** 10^-6^  zval=-4.7030 |
| Fig. S3c | Wilcoxon rank sum | n=22 | p=0.11, 0.15, 3.3x10^-5^, 4.7x10^-7^, 4.7x10^-7^, 4.7x10^-7^  zval=77, 82, 12, 0, 0,0 |
| Fig. S4c | Wilcoxon rank-sum | n=28, sound n=14, context n=5 | compared to safe: shock, sound, ctxt  p=3.0237 **×** 10^-6^, 1.0668 **×** 10^-4^, 9.7055 **×** 10^-4^  zval=4.6692, 3.8749, 3.2989  compared to shock: sound, ctxt  p=0.4422, 0.5277  zval=-0.7685, 0.6315  compared to sound: ctxt  p=0.2282  ranksum= 153.5 |
| Fig. S4d | Wilcoxon rank-sum | U-Maze n=17/17, sound n=10, context n=5 | compared to safe: shock, sound, ctxt  p=0.0136, 2.7893 **×** 10^-5^, 0.0015  zval=-2.4675, -4.1900, -3.1790  compared to shock: sound, ctxt  p=1.2252 **×** 10^-4^, 0.0077  zval=-3.8410, -2.6638  compared to sound: ctxt  p=0.6787  ranksum= 84 |
| Fig. S4i | Spearman correlation | n=33 freezing episodes | R=0.6922  P=8.0848 **×** 10^-6^ |
| Fig. S4f | Wilcoxon signed rank | n=29 | shock  p=0.0495  zval=1.9642  safe  p=0.0112  zval=2.5353 |
| Fig. S5b, Sleep Vs Shock & Sleep Vs Safe | Wilcoxon signed rank | n=24 | compared to sleep: shock, safe  p=0.0072, 0.0177  zval=-2.6857, -2.3724  shock-safe:  p=0.7366  zval=0.3363 |
| Fig. S5c, Sleep Vs Shock & Sleep Vs Safe | Wilcoxon signed rank | n=7 | compared to sleep: shock, safe  p=0.0312, 0.0312  signedrank=1, 1  shock-safe:  p=0.2188  signedrank=22 |
| Fig. S5d, Sleep Vs Shock & Sleep Vs Safe | Wilcoxon signed rank | n=28 | compared to sleep: shock, safe  p=3.7896 **×** 10^-6^, 2.5631 **×** 10^-6^  zval=-4.6226, -4.7030,  shock-safe:  p=0.001  zval=-3.2791 |
| Fig. S5e, Sleep Vs Shock & Sleep Vs Safe | Wilcoxon signed rank | n=19 | compared to sleep: shock, safe  p=1.2042 **×** 10^-4^, 1.6286 **×** 10^-4^  zval=-3.8453, -3.7706  shock-safe:  p=0.0012  zval=3.2303 |
| Fig. S5f, Sleep Vs Shock & Sleep Vs Safe | Wilcoxon signed rank | n=24 | compared to sleep: shock, safe  p=5.2571 **×** 10^-4^, 5.4594 **×** 10^-4^  zval=-3.4673, -3.4571  shock-safe:  p=0.7366  zval=0.3363 |
| Fig. S5g, Sleep Vs Shock & Sleep Vs Safe | Wilcoxon signed rank | n=24 | compared to sleep: shock, safe  p=0.0569, 0.2969  signedrank=3, 7  shock-safe:  p=0.2188  signedrank=6 |
| Fig. S5h, Sleep Vs Shock & Sleep Vs Safe | Wilcoxon signed rank | n=28 | compared to sleep: shock, safe  p=1.7735 **×** 10^-5^, 2.4254 **×** 10^-4^  zval=-4.2917, -3.6700  shock-safe:  p=0.001  zval=-3.2791 |
| Fig. S5i, Sleep Vs Shock & Sleep Vs Safe | Wilcoxon signed rank | n=18 | compared to sleep: shock, safe  p=1.5510 **×** 10^-4^, 3.9814 **×** 10^-4^  zval=-3.7828, -3.5413  shock-safe:  p=0.0012  zval=3.2303 |
| Fig. S6b  Wake, REM, HR, Thigmotaxis | Wilcoxon rank-sum | SDS : n=13,13,13,4*  Ctrl : n= 14,14,8*,3*  **Not all mice were recorded with ECG and video monitoring for thigmotaxis* | p=0.01, 0.0007, 0.93, 0.014  statistic=-2.5, 2.37,27, 93 |
| Fig. S6f  Wake, REM, HR, Thigmotaxis | Wilcoxon rank-sum | Sham : n=10  Rip Inhib: n= 10 | p=0.02, 0.04, 0.02, 0.03  statistic=-2.3, 2 ,46, -2.08 |
| Fig. S6g  Wake, REM, HR, Thigmotaxis | Wilcoxon rank-sum | Saline: n=11  Diazepam: n= 12 | p=0.0095, 0.028, 0.04, 0.35  statistic=2.59, 63, 93, -0.91 |
| Fig. 7a, correlation with safe freq | Spearman correlation | n=29 | R=0.5904  P=0.0018 |
| Fig. S7b, Correlation of parameters with safe side breathing | Pearson/Spearman correlations | n=26, 22, 23, 15, 20 | insomnia, REM sleep, thigmotaxis, HR homecage, HR U-Maze:  R=0.4448, -0.5675, 0.5474, 0.6000, 0.5157  P=0.0238, 0.0067, 0.0077, 0.0199 |
| Fig. S7b, Correlation of parameters with shock side breathing | Pearson/Spearman correlations | n=26, 22, 23, 15, 20 | insomnia, REM sleep, thigmotaxis, HR homecage, HR U-Maze:  R=-0.0869, 0.0593, 0.0999, 0.1929, -0.0271  P=0.6786, 0.7934, 0.6572, 0.4901, 0.9114 |
| Fig. S8a, Correlation SWR density vs breathing rate | Pearson correlation | n=1261 freezing episodes for 29 mice | R=-0.30  P=2.48 **×** 10^-27^ |
| Fig. S8b, Reactivation strength shock versus safe | Wilcoxon signed rank | n=29 | P=0.1120  zval=1.5893 |
| Fig. S8e, vHC stimulation number | Wilcoxon rank-sum | Rip sham=10  Rip inhib=10 | p=0.2123  zval= -1.2473 |
| Fig. S8f, vHC stimulation density in immobility | Wilcoxon rank-sum | Rip sham=10  Rip inhib=10 | shock:  p=0.0889  zval= -1.7015  safe:  p=0.0640  zval= -1.8520 |
| Fig. S8g, SWR disruption effect physiology decoder | Wilcoxon rank-sum | n=10/10 | shock:  P=0.6250  ranksum= 22  safe:  P=0.002  ranksum= 0 |
| Fig. S8h, SWR disruption effect on immobility | Wilcoxon rank-sum | n=10/10 | shock:  P=0.0757  zval=-1.7764  safe:  P=0.9698  zval=-0.0378 |
| Fig. S10a, saline animals | Spearman correlation | n=11 | R=-0.61; P=0.047 |
| Fig. S10b, saline animals vs diazepam animals | Wilcoxon rank-sum | n=11,12 | p=4e-4, statistic=-3.5 |
| Fig. S10c, saline + diazepam animals | Spearman correlation | n=23 | R=-0.64; P=0.0012 |
| Fig. S10d | Wilcoxon rank-sum | n=11/12 | shock:  P=0.0055  ranksum=100  safe:  P=0.3213  ranksum=61 |
| Fig. S10e, effect of DZP on immobility during U-Maze | Wilcoxon rank-sum | n=11/12 | P=0.0455  zval=-2.002 |
| Fig. S10f, effect of DZP on post-conditioning occupancy | Wilcoxon rank-sum | n=11/12 | P=0.8294  zval=-0.2154 |
| Fig. S10g, effect of DZP on SWR occurrence in baseline conditions | Wilcoxon rank-sum | n=11/12 | Quiet wake:  P=0.0235  zval=2.2645  NREM:  P=0.0021  ranksum: 128 |
| Fig. S10h, effect of DZP on immobility in baseline conditions | Wilcoxon rank-sum | n=11/12 | P=0.0513  zval=1.9490 |
| Fig. S10j, effect of DZP on Wake proportion in baseline conditions | Wilcoxon rank-sum | n=11/12 | P=0.8286  ranksum= 98 |
| Fig. S10k, effect of DZP on REM sleep proportion in baseline conditions | Wilcoxon rank-sum | n=11/12 | P=0.5413  zval= 0.6109 |
| Fig. S9a, Saline | Spearman correlation | n=11 | R=0.62; P=0.047 |
| Fig. S9a, Diazepam | Spearman correlation | n=12 | P=0.41 |
| Fig. S9b | Wilcoxon rank-sum | n=11,12 | p=4e-4; statistic=-3.5 |
| Fig. S9c, | Spearman correlation | n=23 | R=0.64; P=0.0012 |
